# An APP-centered molecular gateway integrates innate immunity and retinoic acid signaling to drive irreversible metamorphic commitment

**DOI:** 10.64898/2026.01.22.700939

**Authors:** Ryohei Furukawa, Mizuki Taguchi, Narufumi Kameya, Keisuke Tanaka, Haruka Sato, Takehiko Itoh, Yuh Shiwa

## Abstract

The mechanisms by which environmental signals induce permanent developmental changes remain a fundamental biological problem. We investigated sea star metamorphosis, where microbial biofilms induce a total body plan reorganization. Using systems biology and functional assays, we identified a three-tiered signaling cascade: sensing, conversion, and execution. The immune adaptor MyD88 senses microbes, while MAPK proteins convert this signal into a retinoic acid developmental cue. An amyloid precursor protein (APP)-centered module acts as the irrevocable commitment gateway, stabilized by a positive feedback loop to ensure irreversibility. Remarkably, the genes driving this transition overlap with human pathways for Alzheimer’s disease and ADHD/Autism. By defining this ancient neuro-immune axis in an echinoderm adult body plan similar to the chordate head, our study establishes sea star metamorphosis as a model for understanding the evolutionary origins of human neurological disorders from the evolutionary developmental pathology perspective.

## Introduction

Initiation of metamorphosis, a process involving profound and often irreversible morphological transformations, is a critical stage in the life cycle of many marine invertebrates (*1*, *2*). Crucially, this transition is not autonomous but is triggered by specific microbial signals present in the surrounding environment, primarily those associated with benthic biofilms (*3*, *4*). Widespread reliance on external bacterial cues suggests the presence of a highly conserved molecular system that translates environmental microbial information into complex internal developmental instructions. Understanding the precise molecular link between external microbial recognition and internal commitment to metamorphosis remains a fundamental challenge in evolutionary biology.

In the sea star *Patiria pectinifera*, the decision to settle and undergo metamorphosis is tightly regulated. Previous studies have established that although biofilm is required in nature, metamorphosis can also be chemically induced by exogenous application of retinoic acid (RA) (*5*, *6*). This establishes a functional hierarchy in which environmental recognition ultimately triggers the conserved RA signaling pathway. Additionally, findings from a pharmacological study support this sequence, demonstrating that the RA pathway inhibition halts metamorphosis without affecting the initial larval settlement, indicating that RA signaling operates downstream of environmental recognition, thereby defining its role as a developmental effector within this hierarchical framework (*6*).

Despite this clear functional hierarchy, the specific molecular mechanism that bridges the recognition of the microbial biofilm signal with activation of the internal RA/developmental pathway has remained elusive. Recent studies in the annelid *Hydroides elegans* have identified the innate immune adapter MyD88 as a crucial sensor for microbial signals during metamorphosis (*7*), suggesting an ancient and conserved role for immune signaling in life-cycle transitions. However, whether this MyD88-dependent sensing is conserved in deuterostomes, and how it is functionally converted into the hormonal RA cues that commit developmental fate, remains unknown. This represents a significant barrier to understanding the evolutionary diversification of environmental-developmental signal converters across metazoan phyla.

Besides environmental links, sea star metamorphosis presents a unique opportunity to study the molecular origin of the nervous system. The larval nervous system is systematically destroyed and replaced by a novel, complex adult radial nervous system (*8*). Intriguingly, a recent study suggested that the adult body plan, particularly the organization revealed by anterior-posterior patterning markers, shares striking molecular similarities with the chordate head (forebrain/neural crest territories) (*9*). Therefore, we propose that sea star metamorphosis triggered by environmental cues serves as an evolutionarily accessible model for understanding the ancient molecular mechanisms underlying neural reorganization and patterning, which may contribute to chordate head development.

To resolve the regulatory architecture of this transition in a deuterostome model, we employed a holistic approach combining time-resolved RNA sequencing (RNA-seq), dynamic co-expression network analysis, and comprehensive pharmacological rescue assays. Our results identify an amyloid precursor protein (APP)-centered module as a prime candidate for the integration hub of this transition. Based on our findings, we propose an integrated, multilayered molecular switch model: MyD88 acts as the initial sensor, with a function that is potentially conserved across the protostome-deuterostome divide. At the same time, its signal is functionally converted by JNK/p38 MAPK activity into an RA hormonal cue. We provide evidence suggesting that the APP-centered network structure functions as a critical commitment gateway, accepting input from the RA developmental program and converting it into output for irreversible execution. This study presents a universal mechanistic framework for how environmental microbial signals influence nervous system development, redefining the potential nonpathological role of APP as an ancient environmental-developmental signal converter.

## Results

### Morphological staging of metamorphosis

The precise staging of metamorphosis is crucial for elucidating its molecular basis. To enable an accurate comparison of gene expression during this vital process, we developed a morphological staging system independent of elapsed time. This approach mitigated the significant inter-individual variability observed in the timing of larval settlement and subsequent developmental progression. brachiolaria larvae (Figure 1) were induced to metamorphose using two methods: exposure to biofilm-coated coral sand and treatment with RA.

**Figure 1.**
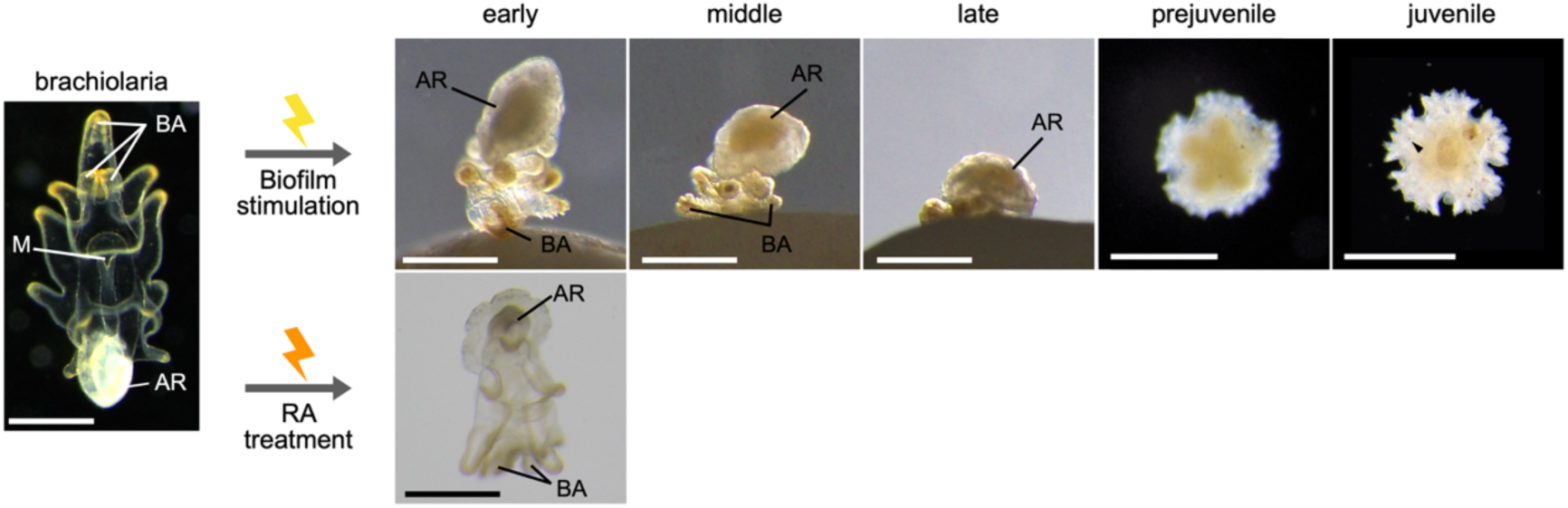
Morphological staging of metamorphosis in *Patiria pectinifera*. The key morphological features of brachiolaria larva before metamorphosis induction are shown: brachiolaria arms (BA), main larval body (M), and adult rudiment (AR). Upon biofilm stimulation (top row), larvae initiate exploratory searching behavior (intermittent attachment/detachment) before settling onto the substrate and progressing irreversibly through the early, middle, late, prejuvenile, and juvenile stages. This progression is characterized by the sequential regression of larval structures and the development of a pentaradial body plan centered on the AR. RA treatment (bottom row) induced morphological transitions comparable to those induced by biofilms, except for initial substrate attachment (settlement). The arrowhead indicates the five ventral red spots that define the juvenile stage. Scale bars, 100 µm.

In coral sand-induced metamorphosis, progression begins with intermittent attachment to and detachment from the substrate via the brachiolaria arms (BA) (exploratory searching behavior). Stable attachment was immediately followed by body inversion, which positioned the adult rudiment (AR) upward. This point was defined as the early stage (Figure 1, biofilm stimulation, early stage). Larval tissue regression was first evident on the aboral side. The middle stage was marked when over half of the larval body had regressed (middle stage). During the late stage, the BA remained visible, whereas the AR expanded into a pentagonal disc (late stage). The subsequent prejuvenile stage was characterized by the complete loss of larval tissues and arms and the emergence of pentaradial symmetry in the AR (prejuvenile stage). Finally, the juvenile stage was defined as the appearance of five red ventral spots aligned along the radial axes (juvenile stage).

RA-induced metamorphosis differed in that the AR expanded into an umbrella-like shape, while the larvae remained swimming. The early stage was defined as concurrent AR expansion and initial larval tissue regression (Figure 1, RA treatment, early stage). However, RA induction required approximately 1.2–1.5 times longer to initiate and exhibited a slower, less synchronous progression than biofilm (bacterial) induction.

Following these morphological criteria, we collected samples from the unstimulated brachiolaria stage and all subsequent metamorphic stages (early, middle, late, prejuvenile, and juvenile) for RNA-seq. Samples were collected from the biofilm induction group (n = 2 per stage, totaling 12 samples from six stages) and the RA induction group (n = 2 only for the early stage). The unstimulated brachiolaria stage was common to both induction groups, resulting in a total of 14 RNA-seq samples. Principal component analysis of the resulting transcriptomic data demonstrated that the samples clustered distinctly by morphological stage (Supplementary Figure 1). Notably, the early-stage samples induced by biofilm and RA formed distinct clusters, positioned between the brachiolaria and middle-stage clusters. This clear separation of the initial response clusters, despite their similar overall positions in the metamorphic trajectory, confirmed that the defined morphological stages accurately reflected major distinct shifts in the underlying transcriptional programs, validating our staging system as a reliable basis for subsequent comparative analysis.

### MyD88-mediated innate immune signaling provides the transcriptional link between biofilm recognition and retinoid signaling

To investigate the molecular link by which external cues initiate metamorphosis, particularly the upstream signaling event that activates the downstream RA pathway, we compared the transcriptomes of early-stage larvae induced by either biofilm stimulation or RA treatment with those of unstimulated larvae. Differential expression analysis identified four major gene sets based on their responses to the two stimuli (biofilm vs. unstimulated and RA vs. unstimulated), including genes that were upregulated or downregulated by one or both stimuli (Figure 2A). For functional analysis, we focused on specific, highly distinct subsets of differentially expressed genes (DEGs) to define stimulus-specific and convergent programs. Differential expression analysis revealed 3,493 upregulated genes and 1,475 downregulated genes under biofilm stimulation, and 1,696 genes were upregulated, and 1,376 were downregulated under RA treatment (Figure 2A).

**Figure 2.**
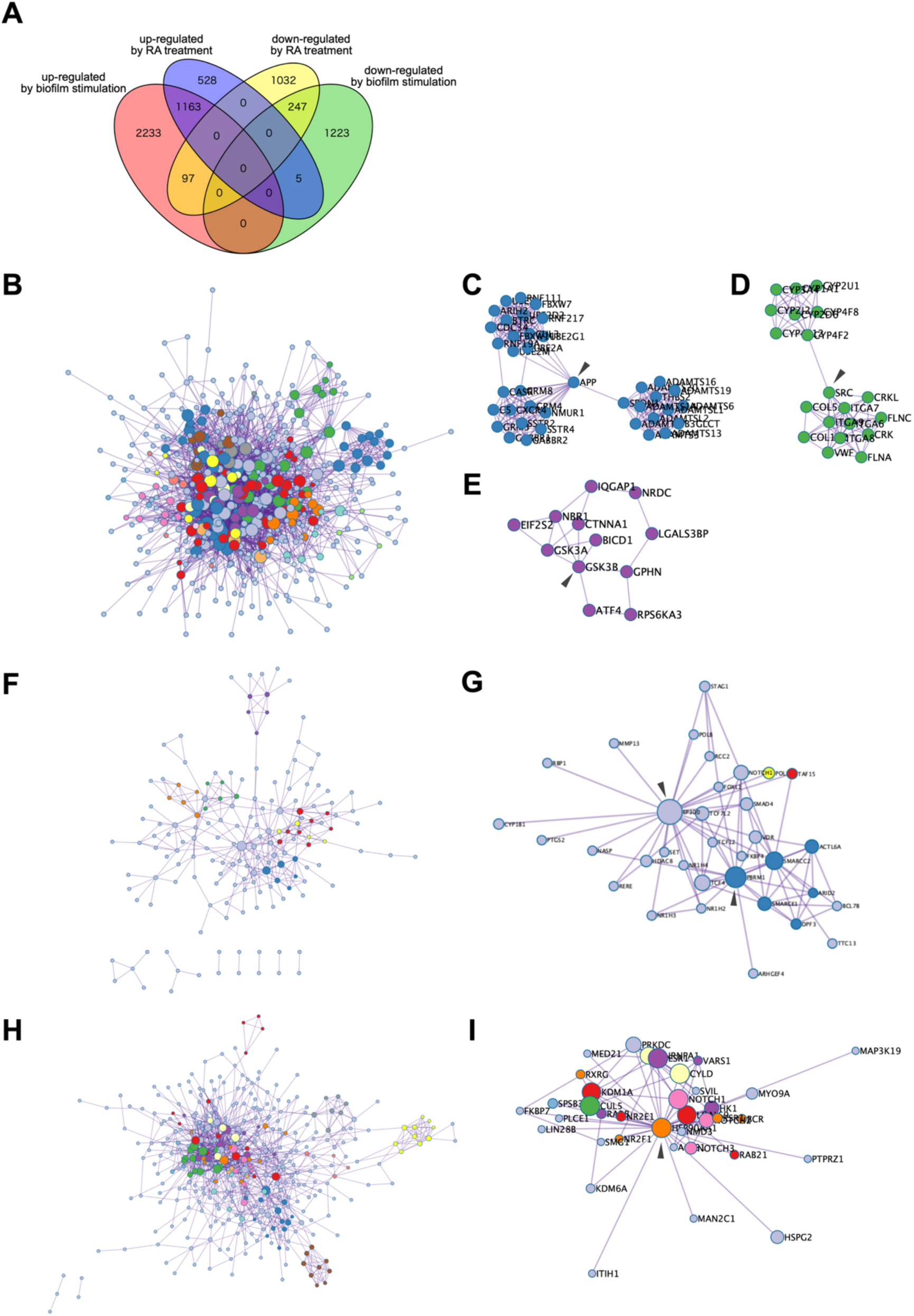
Stimulus-specific and convergent transcriptional programs during early metamorphosis. (A) Venn diagram illustrating the overlap and distinct partitioning of differentially expressed genes (DEGs) at the early stage of metamorphosis, resulting from the comparison of biofilm stimulation (vs unstimulated larva) and retinoic acid (RA) treatment (vs unstimulated larva). The diagram includes both upregulated and downregulated gene sets across the two comparison groups. (B) Protein-protein interaction (PPI) network derived from the biofilm-specific upregulated DEGs (n = 2,233 genes, defined as upregulated by biofilm but not significantly changed by RA). This network is strongly associated with innate immune and MyD88-mediated signaling. (C-E) Subnetworks highlighting central hubs within the biofilm-specific PPI network: (C) APP-, (D) SRC-, and (E) GSK3B-centered subnetworks. (F) PPI network derived from the RA-specific upregulated DEGs (n = 528 genes, defined as upregulated by RA but not significantly changed by biofilm). This network is functionally enriched in developmental and chromatin-related processes. (G) Subnetwork centered on EP300 and PBRM1 within the RA-specific PPI network. (H) Convergent PPI network derived from the upregulated DEGs common to both stimuli (n = 1,163). This network is functionally enriched in MAPK signaling and differentiation pathways. (I) Subnetwork centered on HSP90AA1, which emerges as a major hub integrating the upstream bacterial and RA signaling. In (C-E, G, I), each subnetwork comprises nodes directly connected to the indicated hub protein (black arrowheads).

The biofilm–specific upregulated DEGs (2,233 genes, defined as genes significantly upregulated by biofilm stimulation but not significantly changed by RA treatment) were strongly associated with innate immune processes, prominently “response to bacterium (GO: 0009617),” “MyD88 cascade initiated on plasma membrane (R-HSA-975871),” and “MAPK cascade (GO: 0000165)” (Table 1A). This pronounced enrichment of MyD88 signaling indicates a mechanism by which the bacterial biofilm is initially recognized. Further protein-protein interaction (PPI) network analysis identified APP as a central hub, along with other neurodegenerative disease-related genes, such as SRC and GSK3β (Figure 2B–E). Furthermore, transcription factors, such as NFKB1 (TRR00875), RELA (TRR01158), and RXRA (TRR01177), were predicted to be major transcriptional regulators of these DEGs (Table 1B). Collectively, these findings provide transcriptional evidence that bacterial biofilm signals are rapidly transmitted via a MyD88-dependent innate immune pathway, which likely serves as a crucial molecular link that activates downstream developmental programs.

**Table 1.** Functional and regulatory analysis of stimulus-specific and convergent upregulated.

(A) GO/KEGG enrichment analysis of stimulus-specific and convergent upregulated DEGs.
| GO | Description | Log<sub>10</sub>(P) | Enrichment |
| --- | --- | --- | --- |
| **Biofilm-specific upregulated DEGs** |  |  |  |
| GO:0009617 | response to bacterium | -10.68 | 3.31 |
| R-HSA-975871 | MyD88 cascade initiated on plasma membrane | -10.19 | 5.33 |
| GO:0000165 | MAPK cascade | -9.43 | 2.61 |
| GO:0043065 | positive regulation of apoptotic process | -7.82 | 2.67 |
| R-HSA-112316 | Neuronal System | -7.72 | 3.00 |
| GO:0042063 | gliogenesis | -6.89 | 3.11 |
| GO:0009611 | response to wounding | -6.50 | 2.58 |
| hsa04310 | Wnt signaling pathway | -6.39 | 4.03 |
| hsa04360 | Axon guidance | -6.08 | 3.53 |
| R-HSA-6806834 | Signaling by MET | -5.71 | 4.28 |
| R-HSA-6785807 | Interleukin-4 and Interleukin-13 signaling | -5.16 | 5.29 |
| R-HSA-9006936 | Signaling by TGFB family members | -5.05 | 4.34 |
| GO:0035591 | signaling adaptor activity | -4.92 | 4.23 |
| GO:0031960 | response to corticosteroid | -4.77 | 3.44 |
| M90 | PID WNT CANONICAL PATHWAY | -4.63 | 8.26 |
| GO:0006959 | humoral immune response | -4.52 | 4.20 |
| WP5090 | Complement system in neuronal development and plasticity | -4.39 | 4.09 |
| R-HSA-6794362 | Protein-protein interactions at synapses | -4.39 | 4.09 |
| GO:0043277 | apoptotic cell clearance | -4.37 | 5.50 |
| GO:0045807 | positive regulation of endocytosis | -4.22 | 3.67 |
| GO:0033555 | multicellular organismal response to stress | -3.98 | 4.42 |
| GO:0005112 | Notch binding | -3.98 | 5.67 |
| GO:0043113 | receptor clustering | -3.87 | 4.79 |
| GO:0032526 | response to retinoic acid | -3.66 | 3.72 |
| GO:0050766 | positive regulation of phagocytosis | -3.59 | 5.90 |
| **RA-specific upregulated DEGs** |  |  |  |
| GO:0051048 | negative regulation of secretion | -5.56 | 8.79 |
| GO:0032102 | negative regulation of response to external stimulus | -5.54 | 5.19 |
| GO:0042135 | neurotransmitter catabolic process | -5.50 | 28.00 |
| hsa00830 | Retinol metabolism | -4.40 | 12.42 |
| GO:0004745 | NAD-retinol dehydrogenase activity | -4.38 | 17.50 |
| GO:1902742 | apoptotic process involved in development | -3.65 | 12.70 |
| GO:0016525 | negative regulation of angiogenesis | -3.32 | 7.52 |
| ko04657 | IL-17 signaling pathway | -3.29 | 10.39 |
| WP363 | Wnt signaling pathway | -3.29 | 10.39 |
| GO:0005501 | retinoid binding | -2.88 | 13.18 |
| GO:0070997 | neuron death | -2.81 | 2.71 |
| R-HSA-3371571 | HSF1-dependent transactivation | -2.77 | 11.67 |
| **Convergent upregulated DEGs** |  |  |  |
| R-HSA-1474244 | Extracellular matrix organization | -14.69 | 6.02 |
| GO:0007204 | positive regulation of cytosolic calcium ion concentration | -7.92 | 5.69 |
| GO:0098742 | cell-cell adhesion via plasma-membrane adhesion molecules | -7.42 | 6.13 |
| GO:0004879 | nuclear receptor activity | -7.41 | 13.39 |
| GO:0048598 | embryonic morphogenesis | -6.69 | 3.20 |
| GO:0050808 | synapse organization | -6.40 | 3.54 |
| GO:0000165 | MAPK cascade | -5.10 | 2.70 |
| GO:0050792 | regulation of viral process | -4.83 | 5.27 |
| GO:0032526 | response to retinoic acid | -4.48 | 6.15 |
| WP2849 | Hematopoietic stem cell differentiation | -4.48 | 11.85 |
| R-HSA-114608 | Platelet degranulation | -4.39 | 5.99 |
| GO:0030178 | negative regulation of Wnt signaling pathway | -4.35 | 4.66 |
| GO:2000515 | negative regulation of CD4-positive, alpha-beta T cell activation | -4.03 | 14.23 |
| GO:0072089 | stem cell proliferation | -3.75 | 6.83 |
| GO:0001959 | regulation of cytokine-mediated signaling pathway | -3.70 | 4.84 |
| GO:0008347 | glial cell migration | -3.65 | 6.57 |
| GO:0042501 | serine phosphorylation of STAT protein | -3.39 | 17.07 |
| GO:0007265 | Ras protein signal transduction | -3.12 | 2.13 |
| GO:0002407 | dendritic cell chemotaxis | -2.87 | 12.19 |
| GO:0008037 | cell recognition | -2.45 | 3.09 |
| GO:0042551 | neuron maturation | -2.42 | 5.99 |

(B) Predicted transcriptional regulators identified by TRRUST analysis.
| GO | Description | Log<sub>10</sub>(P) | Enrichment |
| --- | --- | --- | --- |
| **Biofilm-specific upregulated DEGs** |  |  |  |
| TRR00875 | Regulated by: NFKB1 | -4.2 | 2.1 |
| TRR01158 | Regulated by: RELA | -3.6 | 2.0 |
| TRR00344 | Regulated by: FOSL1 | -2.8 | 5.9 |
| TRR00117 | Regulated by: CIITA | -2.4 | 4.9 |
| TRR01177 | Regulated by: RXRA | -2.4 | 4.9 |
| TRR00415 | Regulated by: GATA6 | -2.0 | 5.5 |
| **RA-specific upregulated DEGs** |  |  |  |
| TRR00253 | Regulated by: EGR1 | -3.5 | 5.3 |
| TRR00646 | Regulated by: JUNB | -3.5 | 19.0 |
| TRR01421 | Regulated by: TP63 | -3.5 | 19.0 |
| TRR01521 | Regulated by: VHL | -3.2 | 16.0 |
| TRR00471 | Regulated by: HDAC4 | -3.0 | 13.0 |
| TRR00270 | Regulated by: EP300 | -2.4 | 5.9 |
| TRR00412 | Regulated by: GATA3 | -2.4 | 8.5 |
| TRR00645 | Regulated by: JUN | -2.3 | 3.8 |
| TRR01256 | Regulated by: SP1 | -2.3 | 2.2 |
| TRR00108 | Regulated by: CDX2 | -2.0 | 6.7 |
| TRR00705 | Regulated by: LEF1 | -2.0 | 6.7 |
| **Convergent upregulated DEGs** |  |  |  |
| TRR00263 | Regulated by: ELK1 | -3.2 | 8.1 |
| TRR01158 | Regulated by: RELA | -2.9 | 2.3 |
| TRR01225 | Regulated by: SIRT1 | -2.4 | 4.5 |
| TRR00875 | Regulated by: NFKB1 | -2.3 | 2.1 |
| TRR00351 | Regulated by: FOXD3 | -2.2 | 7.1 |
| TRR00410 | Regulated by: GATA1 | -2.2 | 3.4 |
| TRR01256 | Regulated by: SP1 | -2.1 | 1.7 |
| TRR00125 | Regulated by: CREB1 | -2.0 | 3.2 |

In contrast, RA-specific upregulated DEGs (528 genes, defined as genes significantly upregulated by RA treatment but not significantly altered by biofilm stimulation) were highly enriched in developmental and epigenetic processes, specifically in pathways related to “Retinol metabolism (hsa00830),” “apoptotic process involved in development (GO:1902742),” and “Wnt signaling (WP363)” (Table 1A). The PPI network revealed hub genes, such as the chromatin remodeling factor EP300 and the immunomodulatory factor PBRM1 (Figure 2F and G), with predicted regulators, including HDAC4 (TRR00471) and other epigenetic modifiers (Table 1B). These results confirm that RA stimulation directly activates transcriptional programs associated with retinoid metabolism and chromatin regulation.

The convergent upregulated DEGs (1,163 genes, defined as genes upregulated by both stimuli) revealed convergence on pathways related to “synapse organization (GO:0050808),” “MAPK cascade (GO:0000165),” and “hematopoietic stem cell differentiation (WP2849)” (Table 1A). Within the PPI network of these common DEGs, HSP90AA1 emerged as a major hub (Figure 2H and I), and TRRUST analysis predicted shared regulators including RELA (TRR01158), SIRT1 (TRR01225), and NFKB1 (TRR00875) (Table 1B). These shared regulatory factors likely represent critical nodes that integrate upstream bacterial (MyD88-mediated) and RA inputs, thereby providing a molecular gateway linking external cues to the conserved core program of metamorphosis.

### MyD88 signaling functions as a molecular trigger for metamorphosis initiation

RNA-seq analysis revealed that biofilm stimulation upregulated multiple innate immune genes, particularly those associated with MyD88-dependent signaling, suggesting that this pathway may act as the initial trigger of metamorphosis. To functionally validate this and define the signaling hierarchy, we applied specific pharmacological inhibitors and assessed their effects on larval attachment/settlement and subsequent metamorphic transformation (Figure 3).

**Figure 3.**
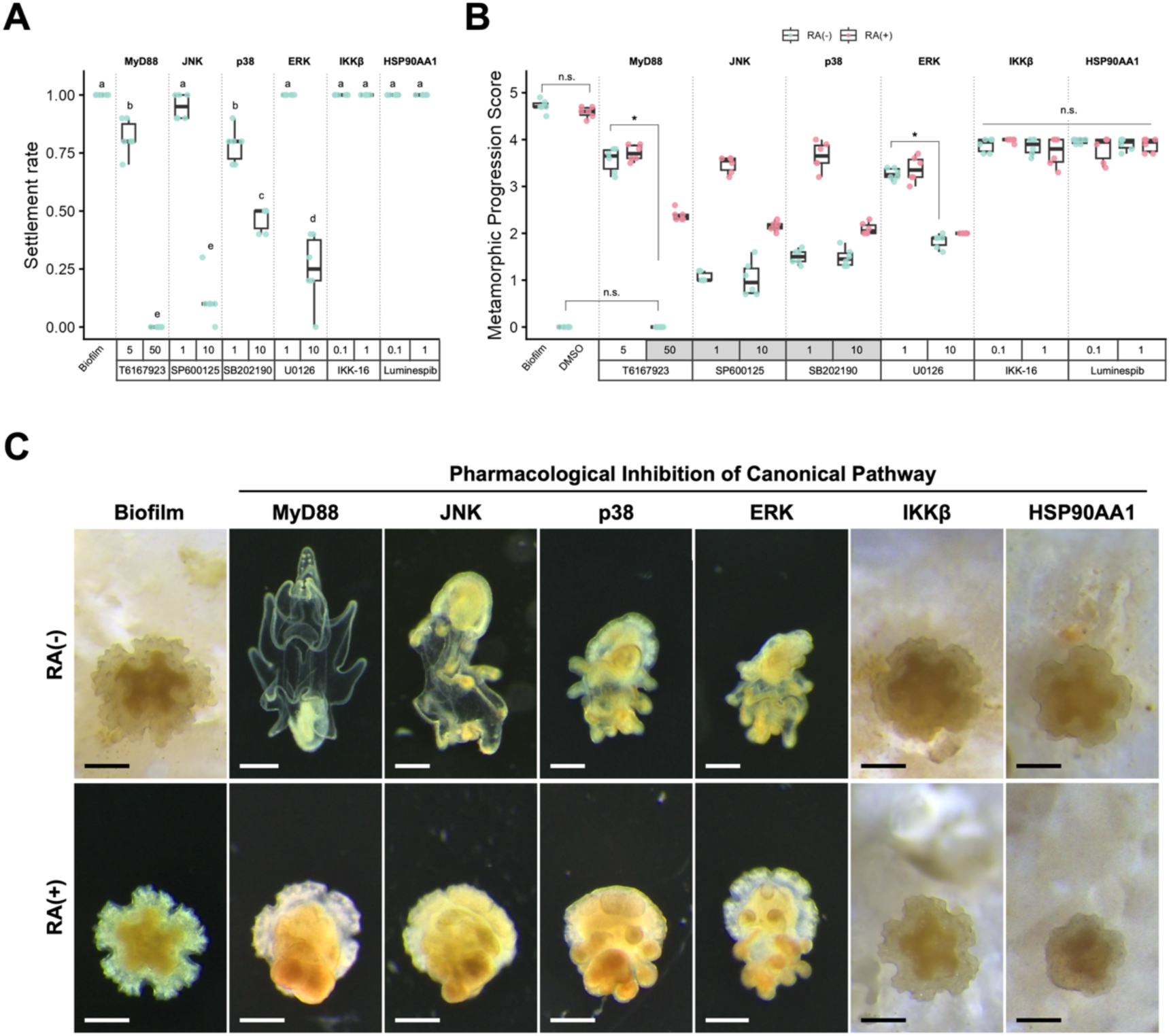
Pharmacological dissection reveals the signaling hierarchy of metamorphosis. (A) Settlement rate of inhibitor-treated larvae. The box plots with superimposed jitter plots display the larval settlement rate under various inhibitor treatments. The biofilm stimulus condition is used as the positive control. The concentration of each inhibitor is indicated on the horizontal axis. The data represent the settlement rate of larvae remaining attached out of 10 larvae across six independent biological replicates (total n = 60). Statistical significance among treatment groups was assessed using one-way ANOVA followed by Tukey’s HSD post hoc test, with grouping letters indicating significant differences (*p* < 0.05); treatments sharing a letter are not significantly different. (B) Quantitative assessment of the functional hierarchy. The box plots with superimposed jitter plots show the Metamorphic Progression Scores (MPS) for larvae treated with various pharmacological inhibitors with or without all-trans retinoic acid (RA). The MPS was calculated based on the metamorphic stage reached by the larvae in the identical assays used for the settlement rate analysis in (A). The concentration of each inhibitor is indicated on the horizontal axis. The MPS represents the average metamorphic stage reached (0 = brachiolaria; 1 = early; 2 = middle; 3 = late; 4 = pre-juvenile; 5 = juvenile). Statistical significance among the treatment groups was assessed using one-way ANOVA followed by Tukey’s HSD post hoc test (*\*p* < 0.05; n.s., not significant). A significant RA-dependent rescue condition (a statistically significant increase in MPS compared with the inhibitor-alone condition) is highlighted in grey, establishing the functional hierarchy of the pathways relative to the RA commitment signal. (C) Representative image illustrating pathway functional hierarchy. Images show representative larval morphology under the control, inhibitor-only, and inhibitor + RA conditions. These images specifically represent the high-concentration inhibitor treatments (MyD88 inhibitor: 50 µM; MAPK inhibitors: 10 µM; IKKβ and HSP90AA1 inhibitors: 1 µM). MyD88 inhibition completely blocks the behavioral decision of settlement. JNK and p38 inhibition caused a distinct early-stage arrest (low MPS), and the effects of their inhibition were significantly rescued by RA co-treatment. In contrast, ERK inhibition arrested metamorphosis at the middle stage, and this block was not rescued by exogenous RA. Similarly, IKKβ and HSP90AA1 inhibition arrested metamorphosis at later stages, and this block was not rescued by exogenous RA, functionally placing all three pathways (ERK, IKKβ, and HSP90AA1) downstream of the RA commitment signal. Scale bar: 200 µm. Inhibitors used: T6167923 (MyD88 inhibitor), IKK-16 (IKKβ inhibitor), U0126 (ERK inhibitor), SP600125 (JNK inhibitor), SB202190 (p38 inhibitor), and Luminespib (HSP90AA1 inhibitor).

To quantitatively evaluate metamorphic progression, we introduced the Metamorphic Progression Score (MPS), which is a weighted average calculated based on the stage reached by individual larvae: brachiolaria (0), early (1), middle (2), late (3), prejuvenile (4), and juvenile (5). RA rescue was defined as a statistically significant increase in MPS in the co-treatment group (inhibitor + RA) compared with that in the inhibitor-only group.

Inhibition of MyD88 by T6167923 suppresses settlement in a concentration-dependent manner. At the high concentration (50 µM), settlement was completely abolished (Settlement rate = 0.00 ± 0.00, MPS = 0.00 ± 0.00; Figure 3A&B). Treatment with exogenous RA did not rescue this inhibited settlement phenotype but significantly rescued metamorphic progression to the middle stage (MPS = 2.38 ± 0.16). This indicates that the MyD88 pathway possesses at least two functionally distinct outputs: one that is essential for the behavioral decision to settle and is not bypassable by RA, and another that triggers the internal molecular program, which is RA-rescuable.

At the lower concentration (5 µM), MyD88 inhibition allowed settlement but significantly reduced maintenance of attachment (Settlement rate = 0.817 ± 0.08), resulting in a late-stage arrest at the prejuvenile stage (MPS = 3.56 ± 0.26). This concentration-dependent shift, from complete settlement block (50 µM) to late-stage molecular arrest (5 µM), suggests that MyD88 signaling also contributes to processes operating downstream of RA commitment, as this late-stage arrest phenotype was not rescued by exogenous RA treatment (MPS = 3.72 ± 0.17).

Crucially, we found that inhibition of the MAPK pathway significantly compromised the ability of larvae to maintain attachment following the initial settlement decision (Figure 3A). High concentrations of JNK (10 µM), ERK (10 µM), and p38 (10 µM) inhibitors resulted in a dramatic and statistically significant loss of attachment (Settlement rates = 0.12 ± 0.10, 0.25 ± 0.15, and 0.47 ± 0.05, respectively). Inhibition of p38 at 1 µM also showed a significant reduction in attachment rate (Settlement rate = 0.78 ± 0.08). This dose-dependent effect, observed primarily in the MAPK pathway inhibitors, indicates that the MyD88-MAPK axis is not only responsible for initial behavioral sensing but also provides an essential signal for the physical stabilization and maintenance of larval attachment.

IKKβ inhibition did not significantly affect the Settlement rate (1.00 ± 0.00) but arrested subsequent metamorphic progression at the pre-juvenile stage (MPS ranging from 3.6 to 4.0). RA co-treatment did not significantly increase MPS (ranging from 3.3 to 4.0) compared with the inhibitor-only group. Similarly, HSP90AA1 inhibition also maintained settlement (Settlement rate = 1.00 ± 0.00), arrested progression at the pre-juvenile stage (MPS ranging from 3.7 to 4.0), and was not rescued by RA co-treatment (MPS ranging from 3.4 to 4.0).

While ERK inhibition drastically reduced attachment at high concentration, inhibition of ERK at low concentration (1 µM) did not prevent attachment (Settlement rate = 1.00 ± 0.00), arrested progression (MPS ranging from 3.1 to 3.4), and was not rescued by RA (MPS ranging from 3.0 to 3.7), regardless of concentration. However, inhibition of JNK and p38 did not significantly prevent attachment but resulted in pronounced arrest at the early stage (MPS ranging from 0.7 to 1.8). RA co-treatment significantly rescued this block; the 1 µM inhibitor concentrations were rescued to the pre-juvenile stage (MPS ranging from 3.2 to 4.0), and the 10 µM concentrations were rescued to the middle stage (MPS ranging from 2.0 to 2.3).

These results establish a functional hierarchy in which JNK and p38 signaling lie upstream of the RA-dependent commitment process, while IKKβ, ERK, and HSP90AA1 function downstream of RA signaling. Furthermore, the data revealed a role for the MyD88-MAPK axis in maintaining the physical stability of post-settled larvae.

### Global co-expression network analysis reveals the integrative regulatory structure of metamorphosis

To gain a comprehensive overview of the transcriptional control architecture throughout the metamorphic process, we performed a gene co-expression network analysis using MEGENA (*10*) (Figure 4A). The DEGs extracted across all metamorphic stages were used to identify highly connected hub genes. Although normalized count data (log-transformed counts per million, log(CPM)) are typically used as inputs for co-expression network construction, the network built using raw count data showed stronger congruence with the early stage comparative data described above, facilitating better biological interpretation than the network built with normalized data (Supplementary Figure 2 and Supplementary Table 2). Therefore, we used a network constructed from the raw count data for all subsequent analyses.

**Figure 4.**
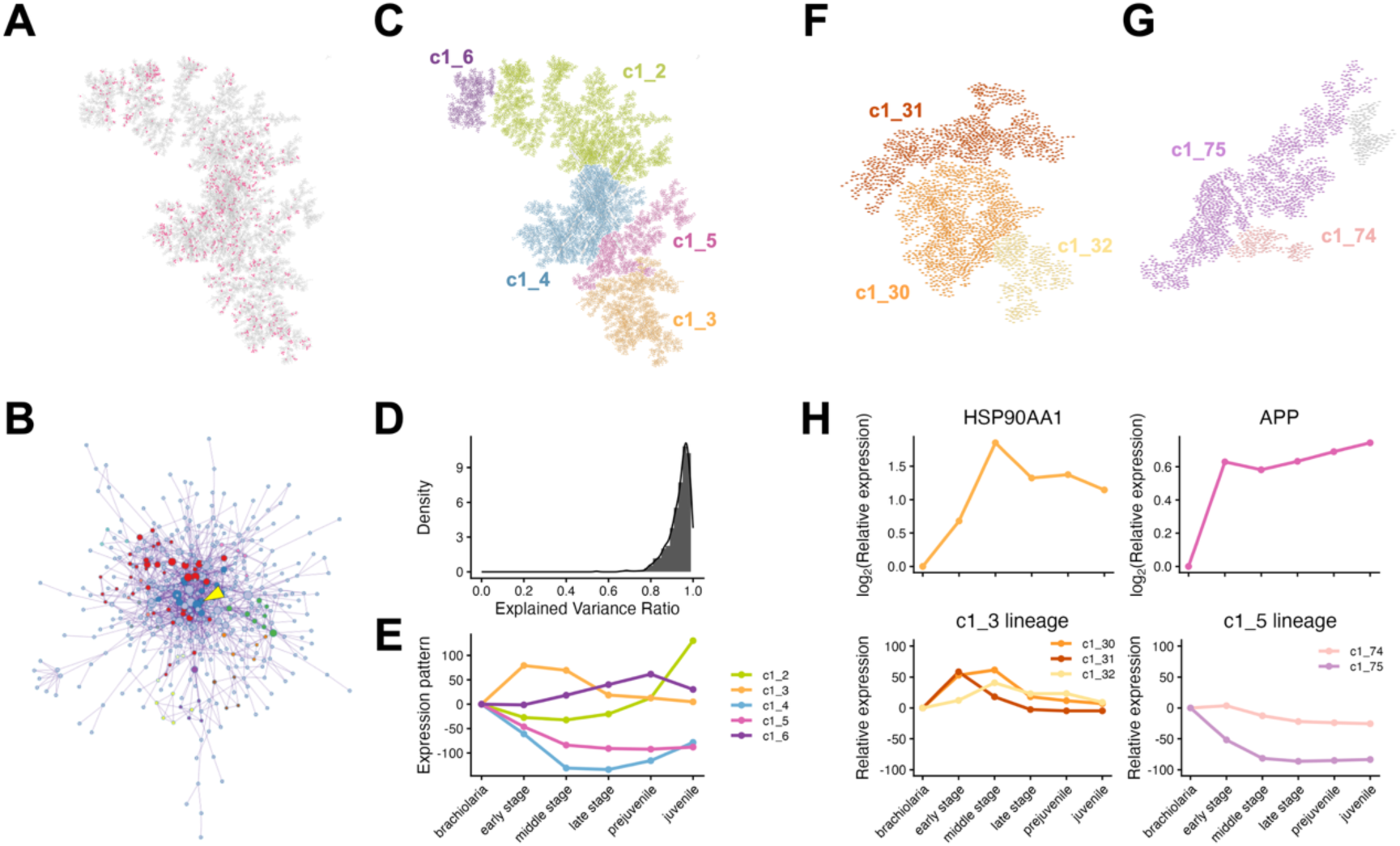
Modular architecture and key hubs of the metamorphic transcriptional network. (A) Global MEGENA network constructed from DEGs across all metamorphic stages. The network reveals the transcriptional control architecture of the transition. Magenta nodes indicate the highly connected hub genes. (B) PPI network of the MEGENA hub genes. HSP90AA1 is highlighted as the node with the highest degree centrality. (C) Modular organization of the MEGENA network, illustrating the five top-layer modules (c1_2 to c1_6). (D) Histogram and density plot showing the distribution of the eigengene’s explanatory power (first principal component). The high variance explained (>80% in most modules) confirms the high accuracy and robust hierarchical structure of the MEGENA construction. (E) Expression trajectories of the top-layer modules (c1_2 to c1_6) across metamorphic stages, represented by their module eigengenes. Each module exhibits a distinct, stage-specific expression pattern. (F, G) Hierarchical sub-modular architecture. (F) The c1_3 network, highlighting its submodules (c1_30, c1_31, c1_32). (G) The c1_5 network, highlighting its submodules (c1_74, c1_75). (H) Comparison of hub gene and module expression patterns. (Top row) Expression profiles of the major hub genes, HSP90AA1 (c1_30 hub, left) and APP (c1_75 hub, right). (Bottom row) Eigengene trajectories of the submodules derived from c1_3 (left) and c1_5 (right). Note that the HSP90AA1 expression profile closely parallels the trends of its parent c1_3 submodules, whereas the APP profile diverges significantly from the overall decreasing trend of the c1_5 lineage, necessitating sub-module analysis.

Functional enrichment analysis of the identified hub genes revealed strong enrichment for crucial biological processes involved in metamorphosis (Table 2). These included “Signaling by Retinoic Acid (R-HSA-5362517),” “tissue remodeling (R-HSA-5362518),” “hematopoietic stem cell differentiation (R-HSA-5362523),” “trans-synaptic signaling (R-HSA-5362528),” “presynapse assembly (R-HSA-5362529),” “response to xenobiotic stimulus (R-HSA-5362531),” and “positive regulation of pattern recognition receptor signaling pathway (GO:0062208)” (Table 2). Furthermore, the PPI network constructed from the hub genes showed that HSP90AA1 had the highest degree centrality (Figure 4B). TRRUST analysis identified TFAP2A (TRR01400) as a major transcriptional regulator in this hub gene cluster (Table 2), indicating that metamorphosis is not driven by a single pathway but rather by the integrated action of a multilayered and multifunctional regulatory network.

**Table 2.** Functional and regulatory analysis of the global MEGENA hub genes.

(A) GO/KEGG enrichment analysis of global hub genes.
| GO | Description | Log<sub>10</sub>(P) | Enrichment |
| --- | --- | --- | --- |
| R-HSA-5362517 | Signaling by Retinoic Acid | -4.14 | 4.42 |
| R-HSA-5362518 | tissue remodeling | -3.80 | 3.06 |
| R-HSA-5362519 | regulation of system process | -3.77 | 1.81 |
| R-HSA-5362520 | positive regulation of muscle contraction | -3.48 | 3.79 |
| R-HSA-5362521 | female pregnancy | -3.40 | 2.41 |
| R-HSA-5362522 | monocarboxylic acid metabolic process | -3.38 | 1.67 |
| R-HSA-5362523 | hematopoietic stem cell differentiation | -3.29 | 6.63 |
| R-HSA-5362524 | mRNA transcription by RNA polymerase II | -3.19 | 4.42 |
| R-HSA-5362525 | secretion | -2.95 | 1.73 |
| R-HSA-5362526 | regulation of hormone levels | -2.93 | 1.63 |
| R-HSA-5362527 | regulation of digestive system process | -2.91 | 4.73 |
| R-HSA-5362528 | trans-synaptic signaling | -2.88 | 1.77 |
| R-HSA-5362529 | presynapse assembly | -2.85 | 3.98 |
| R-HSA-5362530 | axoneme assembly | -2.69 | 1.97 |
| R-HSA-5362531 | response to xenobiotic stimulus | -2.68 | 1.61 |
| GO:1903305 | regulation of regulated secretory pathway | -2.67 | 2.65 |
| GO:0009158 | ribonucleoside monophosphate catabolic process | -2.65 | 5.30 |
| R-HSA-4641263 | Regulation of FZD by ubiquitination | -2.65 | 5.30 |
| GO:0060287 | epithelial cilium movement involved in determination of left/right asymmetry | -2.57 | 3.61 |
| GO:0062208 | positive regulation of pattern recognition receptor signaling pathway | -2.57 | 3.61 |

(B) Predicted transcriptional regulators identified by TRRUST analysis.
| GO | Description | Log<sub>10</sub>(P) | Enrichment |
| --- | --- | --- | --- |
| TRR01400 | Regulated by: TFAP2A | -3.0 | 3.3 |

### Modular architecture of the co-expression network reveals stage-specific transcriptional patterns

To understand the multilayered structure of the gene co-expression network, we divided it into distinct modules (Figure 4C). The eigengene for each module was calculated. An eigengene is defined as the first principal component of the gene expression matrix for each module, representing the typical expression pattern of that module. The explanatory power of the eigengenes for most modules in the MEGENA construction exceeded 80% (Figure 4D). This finding demonstrates that the hierarchical structure of the constructed MEGENA network was built with high accuracy. Five major modules (c1_2–c1_6) were identified in the top layer (Figure 4C). Analysis of expression patterns characterized by eigengenes showed that each top-tier module reached its peak expression at a distinct metamorphic stage (Figure 4E).

The gene HSP90AA1, which we inferred to play a critical role not only in the early stages but also throughout metamorphosis, belongs to the c1_3 module. Furthermore, it was identified as a hub gene in the PPI network, enriched within the c1_3 submodule, c1_30. The expression pattern of HSP90AA1 closely corresponded to that of the c1_3 submodules (Figure 4F and H). Similarly, APP, a hub gene from the early-stage PPI network, was identified as a hub gene in the c1_5 submodule c1_75. In contrast, the expression pattern of APP clearly deviated from that of the parent c1_5 module. Although APP maintained high expression throughout the metamorphic period, the overall expression of the c1_5 module decreased (Figure 4G and H, Supplementary Figure 3). This observed discrepancy between the module-level eigengene and the trajectories of individual hub genes highlights the critical importance of submodule analysis.

### Dynamic network module analysis identifies key regulatory modules driving metamorphic state transitions

Given that the eigengene accurately represented the characteristic expression pattern of its respective module (Figure 4E), each module was treated as a virtual gene. The major strength of MEGENA is its hierarchical modular structure. Leveraging this architecture and eigengenes, we applied the Dynamic Network Biomarker (DNB) theory (*11, 12*) at the module level. The DNB theory captures the coordinated fluctuations of specific gene groups just before the destabilization and transition of a system from one state to another. By applying this to eigengenes, we calculated the Dynamic Network Module (DNM) score as a measure of coordinated fluctuations within modules, thereby identifying the modules that were most strongly associated with the state transition during metamorphosis.

Among the top-layer modules, c1_4 consistently exhibited the lowest DNM scores (Figure 5A, C: c_4 module). Given that the c1_4 eigengene was abruptly downregulated (Figure 4F), this module likely represents the cessation of the larval life program. In contrast, c1_3 and c1_6 showed relatively high scores, c1_2 displayed a bimodal distribution, and c1_5 exhibited a localized peak (Figure 5A). Mapping of DNM scores across all modules revealed that submodules exhibiting high or low DNM scores were heterogeneously distributed within their respective parent modules (Figure 5B).

**Figure 5.**
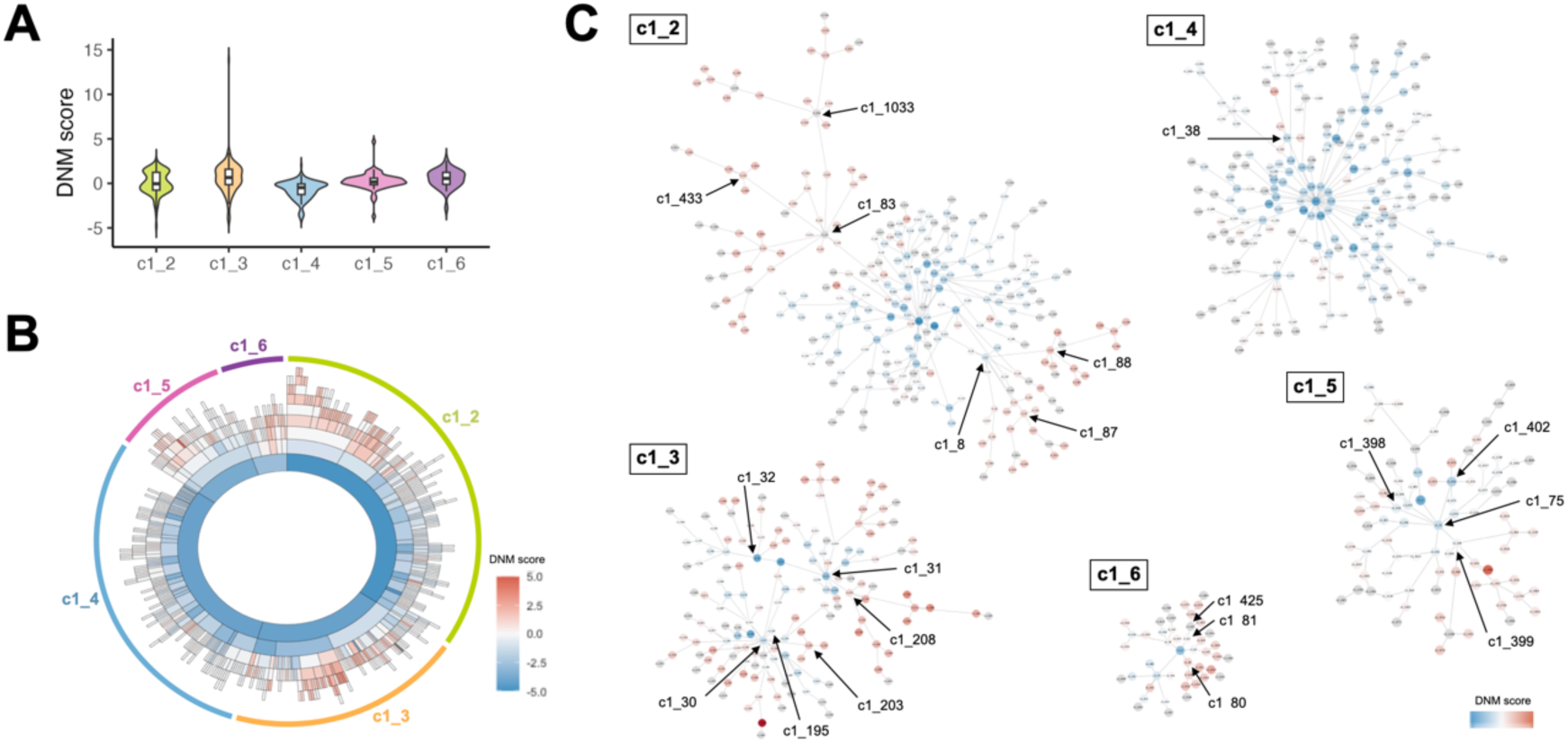
Dynamic network module (DNM) analysis identifies regulatory hubs that drive metamorphic state transitions. (A) Distribution of DNM scores across the five top-layer MEGENA module lineages (c1_2 to c1_6), shown as violin plots. The DNM score quantifies coordinated fluctuations within modules and serves as an indicator of involvement in the developmental state transition. (B) Radial hierarchical map (sunburst plot) visualizing the distribution of DNM scores across all modules in the MEGENA network. Modules are color-scaled by their DNM score (blue: low score/stability; red: high score/fluctuation). The heterogeneous distribution highlights sub-modules undergoing significant coordinated change. (C) Network visualization of module lineages (c1_2 to c1_6). This panel depicts the network structure of each top-layer lineage, with individual nodes colored by their DNM score, as defined in (B). This visualization highlights specific sub-modules that are significantly associated with the state transition. Key sub-modules rich in high DNM scores, such as the APP-centered c1_75 (c1_5 lineage), the HSP90AA1-containing c1_30 (c1_3 lineage), and the SRC-containing c1_83 (c1_2 lineage), are indicated by arrows and module numbers.

Subsequently, we focused on the parent modules that harbored submodules with high DNM scores. The c1_3 module, which was highly expressed in the early to middle metamorphic stages, contained numerous sub-modules with generally high DNM scores (Figure 5B, C: c1_3 module). The c1_31 lineage, which was the earliest to be upregulated within c1_3 (Figure 4I), featured enriched functional annotations related to inflammation, including NFκB signaling, synapse organization, and bacterial response (Table 3). Immune response-related functions were also enriched in c1_30, a submodule containing HSP90AA1 as a PPI network hub. Notably, the c1_30 PPI network also included key hub genes, GSK3β and SIRT1, that were previously detected as hubs in the early-stage metamorphosis PPI network (Figure 2), thereby accurately reflecting the gene expression profile of the initial process. Furthermore, the c1_30 submodule c1_203 showed annotations related to neuronal development and plasticity. c1_32, similar to c1_30, was associated with immune response, alongside terms linked to axon guidance and neuron projection morphogenesis (Table 3).

**Table 3:** Functional and regulatory enrichment of dynamic network submodules.

| Module |  | GO | Description | Log<sub>10</sub>(P) | Enrichment |
| --- | --- | --- | --- | --- | --- |
| c1_2 |  |  |  |  |  |
|  | c1_8 | GO:0099536 | synaptic signaling | -12.40 | 4.55 |
|  |  | GO:0050804 | modulation of chemical synaptic transmission | -6.97 | 3.21 |
|  |  | GO:0006836 | neurotransmitter transport | -5.68 | 4.61 |
|  |  | R-HSA-112316 | Neuronal System | -5.12 | 3.33 |
|  |  | WP5083 | Neuroinflammation and glutamatergic signaling | -3.92 | 4.41 |
|  |  | GO:0051588 | regulation of neurotransmitter transport | -3.32 | 4.67 |
|  |  | GO:0031175 | neuron projection development | -2.79 | 1.95 |
|  |  | GO:1900242 | regulation of synaptic vesicle endocytosis | -2.71 | 10.00 |
|  |  | GO:0140058 | neuron projection arborization | -2.70 | 6.67 |
|  |  | GO:0099170 | postsynaptic modulation of chemical synaptic transmission | -2.53 | 6.06 |
|  |  | R-HSA-8856825 | Cargo recognition for clathrin-mediated endocytosis | -2.11 | 4.76 |
|  | c1_83 | hsa04080 | Neuroactive ligand-receptor interaction | -10.13 | 4.28 |
|  |  | GO:0099536 | synaptic signaling | -8.23 | 3.13 |
|  |  | GO:0010001 | glial cell differentiation | -4.25 | 3.21 |
|  |  | GO:0032228 | regulation of synaptic transmission, GABAergic | -4.19 | 6.89 |
|  |  | GO:0032722 | positive regulation of chemokine production | -3.34 | 6.38 |
|  |  | GO:0009617 | response to bacterium | -3.33 | 2.27 |
|  |  | GO:0014015 | positive regulation of gliogenesis | -3.16 | 4.92 |
|  |  | GO:0099505 | regulation of presynaptic membrane potential | -3.13 | 7.65 |
|  |  | GO:0001764 | neuron migration | -2.99 | 2.87 |
|  |  | GO:0006909 | phagocytosis | -2.85 | 2.94 |
|  |  | GO:0070374 | positive regulation of ERK1 and ERK2 cascade | -2.82 | 3.13 |
|  |  | hsa04145 | Phagosome | -2.30 | 2.87 |
|  |  | GO:0031175 | neuron projection development | -2.24 | 1.65 |
|  |  | GO:0019730 | antimicrobial humoral response | -2.11 | 4.59 |
|  |  | GO:0140058 | neuron projection arborization | -2.11 | 4.59 |
|  | c1_87 | GO:0099536 | synaptic signaling | -5.81 | 4.89 |
|  |  | R-HSA-112316 | Neuronal System | -4.54 | 4.87 |
|  |  | WP5083 | Neuroinflammation and glutamatergic signaling | -3.88 | 7.30 |
|  |  | GO:0006898 | receptor-mediated endocytosis | -3.30 | 5.78 |
|  |  | hsa04726 | Serotonergic synapse | -2.94 | 8.28 |
|  |  | WP5420 | ADHD and autism ASD pathways | -2.43 | 3.12 |
|  |  | GO:0031175 | neuron projection development | -2.30 | 2.42 |
|  |  | GO:0007422 | peripheral nervous system development | -2.00 | 6.54 |
|  | c1_88 | GO:0007268 | chemical synaptic transmission | -8.49 | 10.34 |
|  |  | GO:0099170 | postsynaptic modulation of chemical synaptic transmission | -3.73 | 25.12 |
|  | c1_433 | hsa04080 | Neuroactive ligand-receptor interaction | -10.34 | 29.33 |
|  |  | GO:0007268 | chemical synaptic transmission | -3.29 | 10.02 |
|  | C1_1033 | GO:0099536 | synaptic signaling | -3.92 | 3.30 |
|  |  | GO:0001764 | neuron migration | -3.88 | 5.07 |
|  |  | GO:0060251 | regulation of glial cell proliferation | -3.08 | 13.95 |
|  |  | GO:0099505 | regulation of presynaptic membrane potential | -3.08 | 13.95 |
|  |  | GO:0048667 | cell morphogenesis involved in neuron differentiation | -3.05 | 2.84 |
|  |  | GO:0022010 | central nervous system myelination | -2.85 | 11.96 |
|  |  | GO:0099536 | synaptic signaling | -3.92 | 3.30 |
| **c1_3** |  |  |  |  |  |
|  | c1_30 | hsa05130 | Pathogenic Escherichia coli infection | -3.41 | 3.01 |
|  |  | GO:0006897 | endocytosis | -3.27 | 1.85 |
|  |  | GO:0006959 | humoral immune response | -2.35 | 2.46 |
|  |  | R-HSA-1280218 | Adaptive Immune System | -2.13 | 1.63 |
|  |  | R-HSA-2029482 | Regulation of actin dynamics for phagocytic cup formation | -2.01 | 5.34 |
|  | c1_31 | GO:0043122 | regulation of canonical NF-kappaB signal transduction | -2.82 | 2.74 |
|  |  | GO:0050807 | regulation of synapse organization | -2.71 | 2.25 |
|  |  | R-HSA-1280215 | Cytokine Signaling in Immune system | -2.51 | 1.99 |
|  |  | WP5055 | Burn wound healing | -2.37 | 4.22 |
|  |  | GO:0009617 | response to bacterium | -2.28 | 1.94 |
|  | c1_32 | WP5055 | Burn wound healing | -2.47 | 5.97 |
|  |  | GO:0009617 | response to bacterium | -2.47 | 2.42 |
|  |  | GO:0048812 | neuron projection morphogenesis | -2.31 | 2.15 |
|  |  | R-HSA-422475 | axon guidance | -2.25 | 2.28 |
|  |  | R-HSA-5621481 | C-type lectin receptors | -2.10 | 3.88 |
|  | c1_195 | GO:0006457 | protein folding | -8.40 | 59.81 |
|  |  | C0002726 | Amyloidosis (DisGeNET) | -2.60 | 10.00 |
|  | c1_203 | WP5090 | Complement system in neuronal development and plasticity | -2.71 | 11.70 |
|  |  | GO:0034612 | response to tumor necrosis factor | -2.28 | 8.31 |
|  | c1_208 | hsa04750 | Inflammatory mediator regulation of TRP channels | -3.20 | 9.69 |
|  |  | GO:0001959 | regulation of cytokine-mediated signaling pathway | -2.46 | 6.25 |
|  |  | R-HSA-2142753 | Arachidonate metabolism | -2.13 | 7.27 |
| **c1_4** |  |  |  |  |  |
|  | c1_38 | GO:0030336 | negative regulation of cell migration | -3.14 | 9.42 |
|  |  | GO:2000146 | negative regulation of cell motility | -3.05 | 8.90 |
|  |  | GO:0050680 | negative regulation of epithelial cell proliferation | -2.81 | 12.65 |
|  |  | R-HSA-5684996 | MAPK1/MAPK3 signaling | -2.54 | 10.22 |
|  |  | R-HSA-5673001 | RAF/MAP kinase cascade | -2.54 | 10.22 |
|  |  | R-HSA-5683057 | MAPK family signaling cascades | -2.49 | 9.81 |
| **c1_5** |  |  |  |  |  |
|  | c1_75 | GO:0060055 | angiogenesis involved in wound healing | -4.52 | 7.99 |
|  |  | GO:0001654 | eye development | -3.89 | 2.10 |
|  |  | hsa04360 | Axon guidance | -3.31 | 2.50 |
|  |  | M211 | hedgehog 2 pathway | -3.29 | 5.71 |
|  |  | GO:0009617 | response to bacterium | -3.28 | 2.00 |
|  |  | hsa04148 | Efferocytosis | -2.79 | 2.52 |
|  |  | GO:0006954 | inflammatory response | -2.48 | 1.84 |
|  |  | GO:0051402 | neuron apoptotic process | -2.39 | 2.66 |
|  |  | GO:0007416 | synapse assembly | -2.09 | 2.20 |
|  | c1_398 | GO:0009617 | response to bacterium | -2.51 | 4.92 |
|  | c1_399 | M211 | hedgehog 2 pathway | -5.70 | 17.66 |
|  |  | GO:0001654 | eye development | -3.27 | 3.00 |
|  |  | GO:0009611 | response to wounding | -3.11 | 3.06 |
|  |  | GO:0032526 | response to retinoic acid | -2.55 | 4.94 |
|  | c1_402 | GO:0009617 | response to bacterium | -5.47 | 5.48 |
|  |  | GO:0022010 | central nervous system myelination | -5.23 | 27.34 |
|  |  | GO:0050767 | regulation of neurogenesis | -4.59 | 4.93 |
|  |  | GO:0007422 | peripheral nervous system development | -2.17 | 7.55 |
| **c1_6** |  |  |  |  |  |
|  |  | WP2064 | Neural crest differentiation | -4.23 | 4.19 |
|  |  | GO:0006911 | phagocytosis, engulfment | -2.89 | 6.95 |
|  |  | GO:0002673 | regulation of acute inflammatory response | -2.04 | 6.08 |
|  |  | GO:0043523 | regulation of neuron apoptotic process | -2.02 | 2.11 |
|  |  | GO:0021955 | central nervous system neuron axonogenesis | -2.01 | 3.58 |
|  | c1_80 | GO:0045165 | cell fate commitment | -6.07 | 17.00 |
|  |  | GO:0031175 | neuron projection development | -3.75 | 5.49 |
|  |  | GO:0001708 | cell fate specification | -3.53 | 22.10 |
|  |  | GO:0030900 | forebrain development | -3.08 | 6.46 |
|  | c1_81 | GO:0050768 | negative regulation of neurogenesis | -3.28 | 17.97 |
|  | c1_425 | GO:0050768 | negative regulation of neurogenesis | -3.37 | 19.25 |

The c1_6 lineage, which showed an increased expression after c1_3 (Figure 4E), was primarily annotated by neurogenesis-related processes (Figure 5C: c1_6 module, Table 3). Specifically, the top layer, c1_6, yielded annotations for “Neural crest differentiation (WP2064),” “regulation of acute inflammatory response (GO:0002673),” “regulation of neuron apoptotic process (GO:0043523),” and “central nervous system neuron axonogenesis (GO:0021955).” Its submodules, c1_81 and c1_425, were associated with “negative regulation of neurogenesis (GO:0050768),” while c1_80 included terms for “cell fate commitment (GO:0045165),” “neuron projection development (GO:0031175),” and “forebrain development (GO:0030900).” Taken together, these findings suggest that c1_6 captures fluctuations in neuronal differentiation and specification programs, supporting the transition of neural structures from the larval to the juvenile stage. Following c1_6 expression, the c1_2 lineage, which was upregulated in the juvenile stage (Figure 4E), also demonstrated significant enrichment of neural-associated Gene Ontology (GO) terms in its high-DNM-score sub-modules, including c1_83, which used SRC as the hub of the PPI network. Crucially, c1_2 was not solely restricted to neural development but also showed significant enrichment for innate immune functions, including “response to bacterium (GO:0009617),” “phagocytosis (GO:0006909),” and “antimicrobial humoral response (GO:0019730),” alongside “Neuroinflammation and glutamatergic signaling (WP5083),” and “ADHD and autism ASD pathways (WP5420)” (Figure 5C: c1_2 module, Table 3), suggesting a network heavily centered on continued neural development, integrated with ongoing immune functions at the juvenile stage.

The c1_5 lineage, notably, contained high-DNM-score sub-modules with functional annotations for “response to bacterium (GO:0009617)” (c1_398/c1_402) and “response to retinoic acid (GO:0032526)” (c1_399), and these were found to coexist within the c1_75 module, which harbored APP as its PPI network hub (Figure 5C: c1_5, Table 3). This coexistence in the APP-hub module (c1_75) provides transcriptional evidence that APP directly mediates the signaling bridge between bacterial and RA stimuli, making it the critical signal converter. Additionally, the detection of Hedgehog (Hh) signaling in c1_399 raised the possibility that this pathway cooperates with the RA pathway.

## Discussion

The metamorphic transition of the sea star *Patiria pectinifera* from planktonic larva to benthic juvenile presents a singular model for understanding how environmental signals are translated into irreversible developmental fate decisions. Our study, which combines Dynamic Network Module (DNM) analysis with targeted pharmacological functional assays, resolves the molecular architecture of this process. We established a three-tiered signaling cascade connecting the initial microbial cue to the final neurodevelopmental execution (Figure 6). Recent findings in the annelid *Hydroides elegans* identified

**Figure 6.**
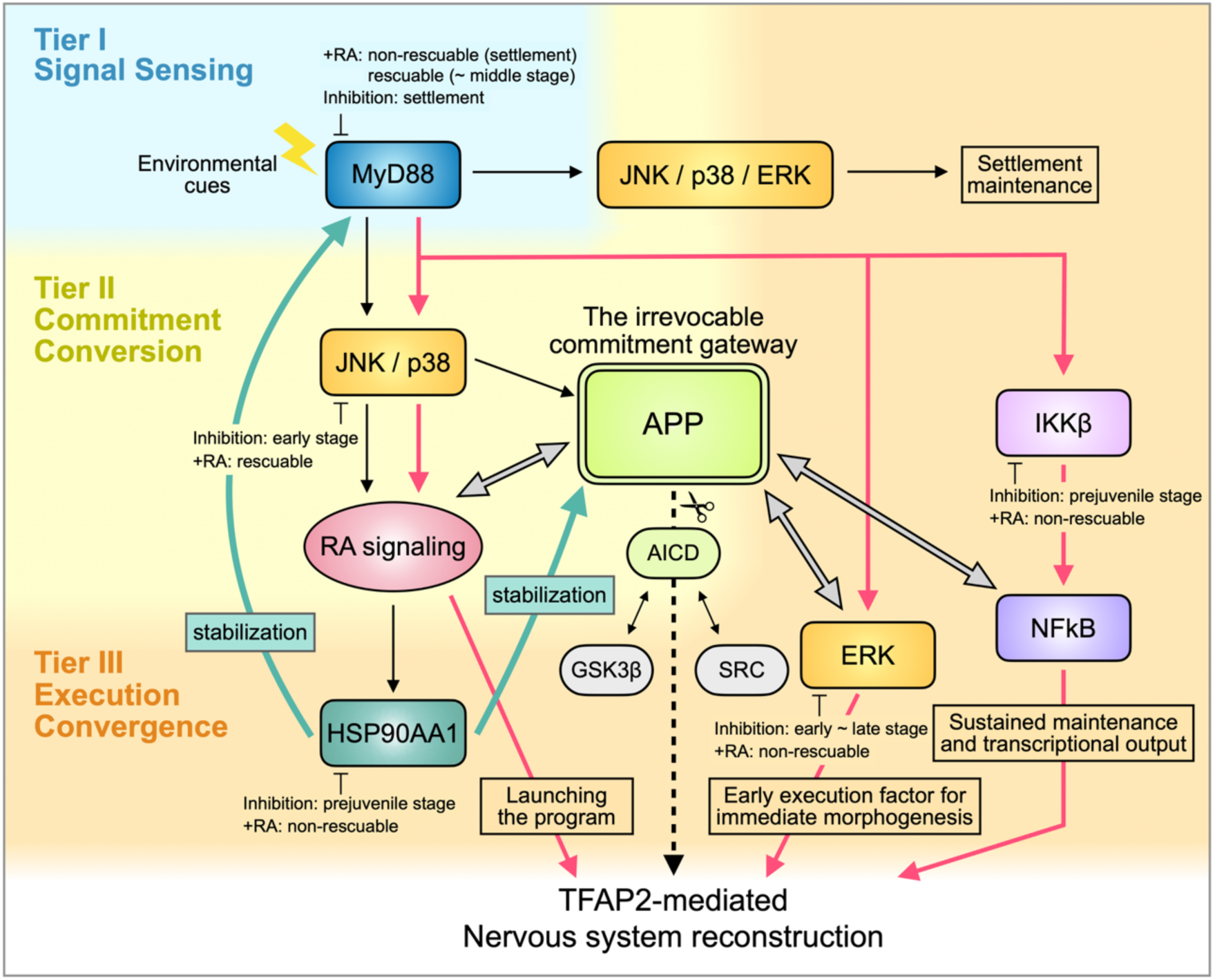
Environmental-Developmental signal converter model for irreversible metamorphic commitment. This model illustrates the proposed three-tiered molecular switch that translates external microbial cues into the irreversible developmental fate of sea star metamorphosis, based on Dynamic Network Module (DNM) analysis and comprehensive pharmacological functional assays. This cascade integrates innate immune and developmental signaling pathways across three functional layers: Signal Sensing, Commitment Conversion, and Irreversible Execution. The process is initiated in the Signal Sensing layer, where the environmental cue, microbial biofilms, activates the adapter protein MyD88, which serves as an obligatory first-tier hub. MyD88 transmits signals via the JNK/p38/ERK MAPK pathway to govern the initial settlement behavior. MyD88 exhibits a concentration-dependent dual output: high-dose inhibition abolishes settlement behavior (RA-non-rescuable), while low-dose inhibition permits settlement but causes a late-stage molecular arrest (RA-non-rescuable). Following sensing, the cascade enters the Commitment Conversion layer. JNK/p38 MAPK acts as an essential hybrid adapter that converts immune signals into a Retinoic Acid (RA) hormonal commitment signal (RA-rescuable phenotype). The Amyloid Precursor Protein (APP) functions as the irrevocable commitment gateway, integrating inputs from the upstream MAPK, IKKβ/NFκB, and RA signaling axes to make the final molecular decision. APP ensures irreversibility through “signal focusing,” maintaining its signal strength during the systemic “mass shutdown” of non-essential larval programs. The process culminates in an Irreversible Execution Tier, where the robust execution of the metamorphic program relies on the multi-layered convergence of signals onto the master transcription factor, TFAP2A. The APP commitment decision is translated into transcriptional output via the release of its intracellular domain (AICD), which acts as the final dedicated execution switch by converging to TFAP2A in complex with GSK3β/Src. TFAP2A receives parallel inputs from RA (for launching the program), IKKβ/NFκB (for sustained maintenance and transcriptional output; RA non-rescuable), and ERK (a crucial early execution factor for immediate morphogenesis and physical attachment maintenance; RA non-rescuable). Finally, the RA signal induces the HSP90AA1 chaperone, establishing a positive feedback loop that maintains the structural integrity and function of critical signaling complexes (including MyD88 and APP), thereby ensuring the stability of the executed program.

MyD88-dependent innate immunity as a key initiator of metamorphosis (*7*). Our results extend this paradigm to the deuterostome lineage, suggesting that the MyD88-MAPK axis is an ancient, evolutionarily conserved “environmental-developmental converter” utilized across diverse metazoan phyla. However, our study reveals that in the sea star, this immune sensing is integrated with an RA-dependent commitment gateway centered on the APP complex, ensuring the finality and irreversibility of the transition.

The metamorphic cascade begins with the recognition of biofilm cues by the MyD88 adapter protein. Pharmacological assays validate the role of MyD88 as an obligatory, first-tier sensor but revealed a concentration-dependent dual-output mechanism. At the high inhibitory concentration (50 µM), MyD88 inhibition completely prevents larval settlement (adhesion), and this behavioral block is not rescued by exogenous RA. However, the internal molecular program was significantly rescued during the middle stage. This demonstrates that MyD88 possesses two functionally distinct outputs: one that serves as the absolute sensor for the behavioral decision to settle (RA-non-rescuable), and another that triggers the internal molecular program (RA-rescuable). Furthermore, at the low concentration (5 µM), MyD88 inhibition permits initial settlement but causes a late-stage arrest at the prejuvenile stage that is RA-non-rescuable. This finding indicates that MyD88 serves not only as an initial trigger but also as a critical maintenance role downstream of the RA commitment process. It likely coordinates with additional execution factors to guarantee the successful completion of the adult body plan.

The signaling hierarchy downstream of MyD88 was precisely defined using RA rescue experiments. While the NFκB branch is canonically regarded as the primary execution axis of MyD88 signaling (*7, 13*), our results in the sea star definitively demonstrate a functional compartmentalization. Specifically, we conclusively show that IKKβ, the core molecule of the NFκB pathway, functions downstream of RA signaling as a maintenance factor. In contrast, the MAPK family members JNK and p38 function as essential “Commitment Converters” located upstream of RA signaling. This RA-rescuable phenotype of JNK/p38 confirms their role in converting immune signals into developmental RA commitment signals, consistent with the known role of MAPK in regulating the enzymes of the retinoid synthesis pathway in developmental contexts (*14–16*). Such regulatory overlap suggests that JNK/p38 triggers RA pathway activation, and provides a necessary temporal brake or fine-tuning mechanism for RA receptor activity, ensuring proper timing of commitment execution.

A functional paradox arises, however, as RA rescue of JNK/p38 inhibition is limited, arresting the program at the pre-juvenile stage (MPS ∼4.0). Our analysis reveals that high concentrations of MAPK inhibitors led to a significant loss of larval attachment (Figure 3A). This demonstrates that JNK/p38 possesses a dual role: it not only serves as an RA signal activator (an RA-rescuable signal conversion function) but also provides the required execution or structural maintenance signal for the physical stability of the settled larva (an RA-non-rescuable function). These findings highlight their function as essential hybrid adaptors in metamorphic processes (*17*). While *Hydroides* co-opted the JNK/p38 axis for direct morphological execution (*18*), the sea star used it primarily as a signal converter for the RA system, demonstrating a subtle but significant evolutionary rewiring of a conserved module.

The components of the downstream “Irreversible Execution Tier” exhibit a RA-non-rescuable phenotype, indicating that they function after the RA commitment signal. This tier is functionally compartmentalized: inhibition of both IKKβ and HSP90AA1 causes a dramatic late-stage developmental arrest, centered around the pre-juvenile stage. This finding places IKKβ in “Execution Maintenance” alongside the chaperone HSP90AA1. Critically, this late-stage arrest temporally correlates with the maximum upregulation of the final execution module c1_2, which is enriched not only for neural development, but also for neuroinflammation and immune response terms. This functional co-localization is highly consistent with the known roles of IKKβ in promoting neural survival and inflammation (*19–21*) and HSP90AA1 in maintaining the structural integrity of critical signaling complexes (*22, 23*).

We propose that the RA commitment signal, driven by JNK/p38, serves as a coordinator that promotes the accumulation of execution factors, specifically HSP90AA1 (c1_30 hub). RA signaling is known to induce the expression of HSP90 family members in response to developmental stress and differentiation (*24*). Crucially, HSP90 is essential for the stability and appropriate processing of APP (*25, 26*). This establishes a positive feedback loop that ensures the structural integrity of the complex signaling machinery–including the MyD88 component and the APP-centered module–during the chaotic process of larval tissue degradation and adult re-patterning. Within this tier, ERK functions as a distinct early execution factor. ERK inhibition results in a later arrest than JNK/p38 but significantly compromises attachment maintenance, suggesting its role in immediate RA-driven morphogenesis and chromatin modulation to facilitate the binding of master regulators like TFAP2A (*27, 28*).

The Amyloid Precursor Protein (APP), identified as the central hub of the c1_75 module, is predicted to act as the irrevocable commitment gateway. Our biofilm-specific PPI network analysis strongly supports this role, identifying APP, GSK3β, and Src as central hubs among DEGs uniquely responsive to microbial cues but independent of RA treatment (Figure 2B-E). This provides transcriptional evidence that the MyD88-dependent signal is channeled into an APP-centered signaling complex.

We propose that APP processing to release its intracellular domain (AICD) serves as the final molecular switch. AICD has been shown to physically interact with GSK3β and Src, forming a transcriptional regulatory complex (*29, 30*). This molecular scaffolding likely facilitates convergence onto the master regulator TFAP2A, which is widely recognized for its crucial roles in neurodevelopment (*27, 31–34*). Crucially, APP maintains high expression throughout metamorphosis, starkly contrasting with the “mass shutdown” of the broader larval program (c1_5 module, Figure 4G). This phenomenon of “signal focusing” represents an active systemic suppression of larval programs and a disconnection from the prior larval architecture. By acting in concert with parallel inputs from RA, MAPK, and IKKβ/NFκB signaling (*19, 35–38*), the APP-driven gateway ensures that the metamorphic stimulus is not only converted but continuously and irreversibly executed, mirroring the robust strategies observed in lineage safeguarding mechanisms (*39, 40*).

Our findings that the metamorphic program is initiated by MyD88-dependent immune response and converted via the MAPK-RA axis into neural reorganization reveal a fundamental evolutionary link between innate immunity and neurodevelopment. Given that the echinoderm adult body plan is molecularly homologous to chordate forebrain and neural crest territories (*9, 41*), our findings define an ancient, non-pathological role for the APP in large-scale neural remodeling (*42–45*). From the perspective of Evolutionary Developmental Pathology (Evo-Devo-Path), human diseases can be viewed as the dysregulation or inappropriate reactivation of such ancestral developmental programs (*46*). The execution module c1_2, enriched for both bacterial response (GO:0009617) and neuroinflammation/glutamatergic signaling (WP5083), embodies this deep evolutionary connection. The role of APP as an “Environmental-Developmental signal converter” in the sea star provides a mechanistic blueprint for understanding how microbial stimuli can trigger pathological neural states in humans. Specifically, clinical links between periodontal pathogens and Alzheimer’s disease (*47–52*) may represent the maladaptive reactivation of this ancient, microbe-responsive neuro-remodeling switch in the aged human brain. Furthermore, the enrichment of ADHD and Autism ASD Pathways within this same execution module suggests that these core programs are evolutionarily ancient and inherently sensitive to regulatory disruption. Within the Evo-Devo-Path framework, the very mechanisms that ensure the robust, rapid construction of a complex nervous system–such as the APP-centered commitment gateway identified here–also represent sites of inherent vulnerability. Sea star metamorphosis thus offers an accessible window into the “Evo-Devo” origins of human “Pathology,” in which the disruption of an ancient neuro-immune regulatory nexus leads to the spectrum of neurodevelopmental and neurodegenerative disorders observed across the metazoan tree.

In conclusion, our integrated analysis delineates a novel three-tiered molecular switch that translates a universal environmental cue into an irreversible metamorphic fate. We functionally defined the MyD88-JNK/p38-RA signaling hierarchy and proposed the APP-centered state transition module, stabilized by an HSP90AA1 feedback loop, as an irreversible commitment gateway. By identifying this ancient neuro-immune bridge, which guides the development of a body plan homologous to the cordate head, we establish sea star metamorphosis as a vital model for Evo-Devo-Path. While direct genetic validation of the APP gateway remains an essential next step, the modular architecture and dynamic commitment points resolved herein provide a robust predictive framework for understanding how the co-option of innate immunity drove the evolution of complex life cycles and the subsequent emergence of human developmental and degenerative diseases.

## Materials and Methods

### Animal maintenance and metamorphosis induction

Adult *Patiria pectinifera* were collected from the coast of Japan and maintained with artificial seawater (MarineArt SF-1, Tomita Pharmaceutical, Japan) at 15 ℃. Embryos were obtained as previously described (*53*). Larvae were reared in ASW at 20 ℃ until they developed into brachiolaria larvae, being fed *Chaetoceros calcitrans* (Sun-culture, Marine Tec, Japan) at a concentration of 1.0 × 10^6^ cells/mL.

Metamorphosis was induced using two methods. For natural induction, brachiolaria larvae were incubated with mature biofilm-coated coral sand (approximately 2–5 mm in diameter) sourced from adult housing tanks (*5*). To ensure the reliability of the natural inducer for all assays, we implemented a biofilm quality control step as described below. For chemical induction, all-trans retinoic acid (RA) (100 µM stock in dimethyl sulfoxide (DMSO); Sigma-Aldrich, US; stored at -80 °C) was added to ASW to a final concentration of 1 µM (*6*). For the chemical induction, the final dimethyl sulfoxide (DMSO) concentration in the carrier control was maintained at 0.1%.

### Morphological staging and phenotypic quantification

Accurate staging of metamorphosis was performed based on morphological criteria, independent of elapsed time, to mitigate interindividual variability (Figure 1). We defined five key metamorphic stages: early, middle, late, pre-juvenile, and juvenile, using distinct, observable external features, such as the extent of larval tissue regression and development of the adult rudiment (AR). This staging system was used for both RNA-seq sample collection and phenotypic observations in the inhibition assays. We calculated the Metamorphic Progression Score (MPS) for phenotypic quantification using pharmacological assays. MPS served as a quantitative measure of the average metamorphic state of the larvae in each experimental group. The MPS was calculated as the weighted average of the number of individuals reaching each stage, with the following weights: brachiolaria (0), early (1), middle (2), late (3), prejuvenile (4), and juvenile (5).

### Pharmacological Inhibition Assays

To functionally validate the signaling pathways identified by RNA-seq analysis, we performed inhibition assays using biofilm-coated coral sand, which reliably induced metamorphosis. All assays were terminated and quantified 48 h post-induction, which was determined to be the time point at which the corresponding DMSO carrier control groups consistently achieved 100% complete metamorphosis (juvenile stage; MPS = 5.0).

Before inhibitor treatment, the biofilm-coated coral sand underwent a quality control (QC) assay. Individual batches of sand were incubated with brachiolaria larvae in DMSO carrier control media for at least 24 h. Only sand batches that successfully induced 100% metamorphosis in the control larvae were selected for the subsequent inhibition assays. This procedure mitigates the variability arising from non-uniform biofilm quality.

The DMSO concentration in the carrier control media varied depending on the inhibitor being tested, ranging from 0.01% to 5% (v/v). Specifically, the DMSO concentration for the MyD88 inhibitor was 0.5% or 5% (corresponding to 5 µM and 50 µM test concentrations); for the JNK, p38, and ERK inhibitors, the concentration was 0.1% or 1.0% (corresponding to 1 µM and 10 µM test concentrations); and for the IKKβ and HSP90AA1 inhibitors, the concentration was 0.1% or 1.0% (corresponding to 0.1 µM and 1 µM test concentrations).

Following QC, the selected sand was incubated in ASW containing chemical inhibitors for the metamorphosis of fresh brachiolaria larvae. The inhibitors were dissolved in DMSO and applied at the indicated concentrations: the MyD88 inhibitor T6167923 (5 or 50 µM; MedChemExpress), the IKKβ inhibitor IKK-16 (0.1 or 1 µM; MedChemExpress), and MAPK inhibitors SP600125 (JNK), SB202190 (p38), and U0126 (ERK) (1 or 10 µM; MedChemExpress or FUJIFILM Wako Pure Chemical Corporation), and the HSP90AA1 inhibitors luminespib (0.1 or 1 µM; Chemscene). The final DMSO concentration under each inhibitor condition was matched to the corresponding carrier control concentration as described above.

To test whether the inhibited pathways acted upstream of RA signaling, parallel experiments were conducted in which all-trans RA was co-administered with the inhibitors. The RA was added to the induction medium at its full effective concentration (1 µM) alongside the inhibitors. The inhibitor + RA conditions were run concurrently with the respective DMSO carrier controls.

The larvae were then incubated with various inhibitors for the entire duration of 48 h. At least three independent biological replicates were used for each condition (n = 10 larvae per group). Morphological progression was quantified by measuring MPS at the 48-hour final time point. The definition of RA rescue as a statistically significant increase in MPS in the co-treatment group relative to the inhibitor-only group is detailed in the Results Section.

### Genome sequencing and assembly

Genomic DNA was extracted from the testis tissue of *Patiria pectinifera* collected at the Misaki Marine Biological Station, using standard protocols. DNA quality and quantity were assessed using a Qubit Fluorometer and a NanoDrop spectrophotometer. Illumina paired-end libraries with an insert size of approximately 500 bp were prepared using the TruSeq DNA Sample Preparation Kit (Illumina). Mate-pair libraries with insert sizes of approximately 3 kb, 6 kb, 10 kb, and 15 kb were constructed using the Nextera Mate Pair Sample Preparation Kit (Illumina). Sequencing was performed on an Illumina HiSeq 2500 platform, generating 2 × 250 bp paired-end reads for paired-end libraries and 2 × 100 bp paired-end reads for mate-pair libraries. Genome assembly was performed using Platanus-allee v2.2.1 (*54*), followed by scaffolding with mate-pair libraries and gap closing using paired-end reads.

### RNA-seq sample preparation and data acquisition

RNA-seq was performed on brachiolaria larvae and five metamorphic stages. Samples for the metamorphic stages were collected either by carefully detaching them from the coral sand or by following RA-induced metamorphosis. Two independent biological replicates (n = 2) were collected for each condition and stage. Lysates from 20 to 30 individuals were pooled for each biological replicate to ensure sufficient biological variation capture within each sample and achieve a high starting quantity of total RNA (≥ 250 ng). Total RNA was extracted using a NucleoSpin RNA kit (TAKARA). RNA quality and quantity were assessed using a Qubit Fluorometer (Thermo Fisher Scientific), NanoDrop spectrophotometer (Thermo Fisher Scientific), and TapeStation system (Agilent Technologies).

Libraries were prepared from 100 ng total RNA using the NEBNext Ultra II RNA Library Prep Kit for Illumina (New England Biolabs). Sequencing was performed on a NextSeq 500 system (Illumina) using the NextSeq 500/550 Mid Output Kit v2.5 (150 cycles), yielding 2–75 bp paired-end reads.

Sequencing reads were mapped to the *Patiria pectinifera* draft genome using STAR (v2.7.3a). Strand-specific transcriptome assembly was conducted using Scallop (v0.10.4), and functional annotations for the assembled transcriptome were obtained using Trinotate (v3.1.0). Gene expression levels were quantified using featureCounts (v1.5.2), and lowly expressed genes were filtered using the filterByExpr function of the edgeR package (v3.24.3) in R (v3.4.0). Given the limited number of replicates (n = 2), DEGs were identified using the edgeR package, known for its robust performance with low replicate numbers, based on a false discovery rate (FDR)-adjusted P-value (q-value) < 0.05.

### Gene co-expression networks analysis by MEGENA

To construct the gene co-expression networks, we employed a Multiscale Embedded Gene Co-Expression Network Analysis (MEGENA) framework (*10*) (MEGENA R package v1.3.7). The DEGs identified across the metamorphic time course were used. Two types of input data were tested using MEGENA: raw data and normalized counts (log-transformed counts per million, log(CPM)). Because of the limited sample size (n = 2), we opted for raw count data as the input for network construction, as networks derived from raw counts exhibited superior biological interpretability and stronger topological correspondence with the functional enrichment of early-stage DEGs (as detailed in Table 1 and Supplementary Table 1). For each module, the eigengene was computed as the first principal component of the scaled expression profile.

### Dynamic network module (DNM) analysis

To detect early warning signals of critical transitions, we adapted a framework of dynamic network biomarkers (DNBs) (*11, 12*) at the module level. The DNB theory was applied to the MEGENA modules’ eigengenes. This approach minimizes the noise associated with individual gene expression and leverages the stability of module-level signals (eigengenes, which explained >80% of variance; Figure 4D) to overcome the limitation of low replicate number. A composite DNM score was defined as the integration of (i) fluctuation amplitude (ΔCV), (ii) reduction of internal correlations, and (iii) divergence from external modules. Modules exhibiting high DNM scores were considered candidate modules for signaling critical states preceding metamorphic transitions. The detailed procedures are described in the Supplementary Methods section, and the complete R scripts used to implement the workflow are provided in the Supplementary File (DNM_Analysis_files. zip).

### Statistical analysis and functional enrichment

Functional enrichment analyses, including Gene Ontology (GO) terms, KEGG pathways, Protein-Protein Interaction (PPI) networks, and transcriptional regulator analysis (TRRUST), were performed exclusively using the online tool Metascape (https://metascape.org) (*55*) under default settings. Metascape built-in analysis tools were used to identify hub genes within the PPI network and predict upstream transcriptional regulators.

Basic statistical tests for phenotypic quantification were performed using the R software (v4.0.5). To assess the effects of inhibitor treatment on larval settlement and metamorphosis, percentage data were compared using Fisher’s exact test. Additional statistical comparisons among the treatment groups were performed using one-way analysis of variance (ANOVA), followed by Tukey’s HSD post-hoc test. *P*-values less than 0.05 were considered statistically significant.

## Acknowledgments

We thank the members of the Research Center for Marine Biology, Graduate School of Life Sciences, Tohoku University, for supplying *Patiria pectinifera*. We are particularly grateful to Dr. Ritsu Kuraishi and Dr. Midori Matsumoto for their critical discussions.

## Funding

JSPS KAKENHI Grant Number JP23K05400 (RF)

Cooperative Research Grant of the Genome Research for BioResource, NODAI

Genome Research Center, Tokyo University of Agriculture (RF)

Keio University Academic Development Funds (RF)

## Author contributions

Conceptualization: RF, MT

Methodology: RF, MT, KT, HS, TI, YS

Investigation: RF, MT, NK, KT, HS, TI

Formal Analysis: RF, MT, TI

Resources: TI, YS

Visualization: RF, MT

Supervision: RF

Writing—original draft: RF, MT

Writing—review & editing: RF, MT, NK, KT, HS, TI, YS

## Data and materials availability

The draft genome of *P. pectinifera*, along with genomic and functional annotations, has been deposited in Zenodo, an open-access data repository (DOI: 10.5281/zenodo.18250764). Transcriptome datasets have been deposited in the NCBI Sequence Read Archive (SRA) and are available in the International Nucleotide Sequence Database Collaboration databases (including DDBJ, NCBI, and EBI) under the BioProject Accession Number PRJNA1371075. All data required to evaluate the conclusions of this study are presented in the paper and/or Supplementary Materials. The count data used in this article, with low-count reads removed, has been uploaded as the supplementary data (TMM_count.txt).

## References

1. M. G. Hadfield, E. J. Carpizo-Ituarte, K. Del Carmen, B. T. Nedved, Metamorphic competence, a major adaptive convergence in marine invertebrate larvae. Am. Zool. 41, 1123–1131 (2001).

2. M. G. Hadfield, Why and how marine-invertebrate larvae metamorphose so fast. Semin. Cell Dev. Biol. 11, 437–443 (2000).

3. M. G. Hadfield, Biofilms and marine invertebrate larvae: What bacteria produce that larvae use to choose settlement sites. Annu. Rev. Mar. Sci. 3, 453–470 (2011).

4. M. Rischer, H. Guo, C. Beemelmanns, Signalling molecules inducing metamorphosis in marine organisms. Nat. Prod. Rep. 39, 1833–1855 (2022).

5. N. Murabe, H. Hatoyama, M. Komatsu, H. Kaneko, Y. Nakajima, Adhesive papillae on the brachiolar arms of brachiolaria larvae in two starfishes, *Asterina pectinifera* and *Asterias amurensis*, are sensors for metamorphic inducing factor(s). Dev. Growth Differ. 49, 647–656 (2007).

6. S. Yamakawa, Y. Morino, M. Honda, H. Wada, The role of retinoic acid signaling in starfish metamorphosis. EvoDevo. 9, 10 (2018).

7. E. Darin, M. V. Farrell, T. N. Ali, J. Rivera Alfaro, K. E. Malter, N. J. Shikuma, MyD88 knockdown by RNAi prevents bacterial stimulation of tubeworm metamorphosis. Proc. Natl. Acad. Sci. U. S. A. 112, e2505805122 (2025).

8. L. Elia, P. Selvakumaraswamy, M. Byrne, Nervous system development in feeding and nonfeeding asteroid larvae and the early juvenile. Biol. Bull. 216, 322–334 (2009).

9. L. Formery, P. Peluso, I. Kohnle, J. Malnick, J. R. Thompson, M. Pitel, K. R. Uhlinger, D. S. Rokhsar, D. R. Rank, C. J. Lowe, Molecular evidence of anteroposterior patterning in adult echinoderms. Nature. 623, 555–561 (2023).

10. W.-M. Song, B. Zhang, Multiscale embedded gene co-expression network analysis. PLOS Comput. Biol. 11, e1004574 (2015).

11. L. Chen, R. Liu, Z.-P. Liu, M. Li, K. Aihara, Detecting early-warning signals for sudden deterioration of complex diseases by dynamical network biomarkers. Sci. Rep. 2, 342 (2012).

12. K. Aihara, R. Liu, K. Koizumi, X. Liu, L. Chen, Dynamical network biomarkers: Theory and applications. Gene. 808, 145997 (2022).

13. T. Kawai, M. Ikegawa, D. Ori, S. Akira, Decoding toll-like receptors: Recent insights and perspectives in innate immunity. Immunity. 57, 649–673 (2024).

14. Y. Ohoka, A. Yokota-Nakatsuma, N. Maeda, H. Takeuchi, M. Iwata, Retinoic acid and GM-CSF coordinately induce retinal dehydrogenase 2 (RALDH2) expression through cooperation between the RAR/RXR complex and Sp1 in dendritic cells. PLOS One. 9, e96512 (2014).

15. Z. Al Tanoury, S. Gaouar, A. Piskunov, T. Ye, S. Urban, B. Jost, C. Keime, I. Davidson, A. Dierich, C. Rochette-Egly, Phosphorylation of the retinoic acid receptor RARγ2 is crucial for the neuronal differentiation of mouse embryonic stem cells. J. Cell Sci. 127, 2095–2105 (2014).

16. Z. Chai, L. Yang, B. Yu, Q. He, W. I. Li, R. Zhou, T. Zhang, X. Zheng, J. Xie, p38 mitogen-activated protein kinase-dependent regulation of SRC-3 and involvement in retinoic acid receptor α signaling in embryonic cortical neurons. IUBMB Life. 61, 670–678 (2009).

17. S. Wang, H. Li, T. Chen, H. Zhou, W. Zhang, N. Lin, X. Yu, Y. Lou, B. Li, E. Yinwang, Z. Wang, K. Wang, Y. Xue, H. Qu, P. Lin, H. Sun, W. Teng, H. Mou, X. Chai, Z. Cai, Z. Ye, Human γδ T cells induce CD8^+^ T cell antitumor responses via antigen-presenting effect through HSP90-MyD88-mediated activation of JNK. Cancer Immunol. Immunother. 72, 1803–1821 (2023).

18. N. J. Shikuma, I. Antoshechkin, J. M. Medeiros, M. Pilhofer, D. K. Newman, Stepwise metamorphosis of the tubeworm *Hydroides elegans* is mediated by a bacterial inducer and MAPK signaling. Proc. Natl. Acad. Sci. U. S. A. 113, 10097–10102 (2016).

19. B. S. Sivamaruthi, N. Raghani, M. Chorawala, S. Bhattacharya, B. G. Prajapati, G. M. Elossaily, C. Chaiyasut, NF-κB pathway and its inhibitors: A promising frontier in the management of Alzheimer’s disease. Biomedicines. 11, 2587 (2023).

20. I. I. Babkina, S. P. Sergeeva, L. R. Gorbacheva, The role of NF-κB in neuroinflammation. Neurochem. J. 15, 114–128 (2021).

21. M. Mettang, S. N. Reichel, M. Lattke, A. Palmer, A. Abaei, V. Rasche, M. Huber-Lang, B. Baumann, T. Wirth, IKK2/NF-κB signaling protects neurons after traumatic brain injury. FASEB J. 32, 1916–1932 (2018).

22. M. Hinz, C. Scheidereit, The IκB kinase complex in NF-κB regulation and beyond. EMBO Rep. 15, 46–61 (2014).

23. A. Israël, The IKK complex, a central regulator of NF-κB activation. Cold Spring Harb. Perspect. Biol. 2, a000158 (2010).

24. N. L. Nguyen, T. X. Hoang, J. Y. Kim, All-trans retinoic acid-induced cell surface heat shock protein 90 mediates Tau protein internalization and degradation in human microglia. Mol. Neurobiol. 62, 742–755 (2025).

25. A. A. Noorani, H. Yamashita, Y. Gao, S. Islam, Y. Sun, T. Nakamura, H. Enomoto, K. Zou, M. Michikawa, High temperature promotes amyloid β-protein production and γ-secretase complex formation via Hsp90. J. Biol. Chem. 295, 18010–18022 (2020).

26. R. E. Lackie, A. Maciejewski, V. G. Ostapchenko, J. Marques-Lopes, W.-Y. Choy, M. L. Duennwald, V. F. Prado, M. A. M. Prado, The Hsp70/Hsp90 chaperone machinery in neurodegenerative diseases. Front. Neurosci. 11, 254 (2017).

27. C. Jin, Y. Luo, Z. Liang, X. Li, D. Kołat, L. Zhao, W. Xiong, Crucial role of the transcription factors family activator protein 2 in cancer: Current clue and views. J. Transl. Med. 21, 371 (2023).

28. C. I. Semprich, L. Davidson, A. Amorim Torres, H. Patel, J. Briscoe, V. Metzis, K. G. Storey, ERK1/2 signalling dynamics promote neural differentiation by regulating chromatin accessibility and the polycomb repressive complex. PLOS Biol. 20, e3000221 (2022).

29. F. Zhou, K. Gong, B. Song, T. Ma, T. van Laar, Y. Gong, L. Zhang, The APP intracellular domain (AICD) inhibits Wnt signalling and promotes neurite outgrowth. Biochim. Biophys. Acta, Mol. Cell Res. 1823, 1233–1241 (2012).

30. H. Bukhari, A. Glotzbach, K. Kolbe, G. Leonhardt, C. Loosse, T. Müller, Small things matter: Implications of APP intracellular domain AICD nuclear signaling in the progression and pathogenesis of Alzheimer’s disease. Prog. Neurobiol. 156, 189–213 (2017).

31. H. Kantarci, R. K. Edlund, A. K. Groves, B. B. Riley, Tfap2a promotes specification and maturation of neurons in the inner ear through modulation of Bmp, Fgf and notch signaling. PLOS Genet. 11, e1005037 (2015).

32. N. de Crozé, F. Maczkowiak, A. H. Monsoro-Burq, Reiterative Ap2A activity controls sequential steps in the neural crest gene regulatory network. Proc. Natl. Acad. Sci. U. S. A. 108, 155–160 (2011).

33. J. Holzschuh, A. Barrallo-Gimeno, A.-K. Ettl, K. Dürr, E. W. Knapik, W. Driever, Noradrenergic neurons in the zebrafish hindbrain are induced by retinoic acid and require *tfap2a* for expression of the neurotransmitter phenotype. Development. 130, 5741–5754 (2003).

34. K. Yang, J. Zhao, S. Liu, S. Man, RELA promotes the progression of oral squamous cell carcinoma via TFAP2A-Wnt/β-catenin signaling. Mol. Carcinog. 62, 641–651 (2023).

35. G. König, C. L. Masters, K. Beyreuther, Retinoic acid induced differentiated neuroblastoma cells show increased expression of the *β*A4 amyloid gene of Alzheimer’s disease and an altered splicing pattern. FEBS Lett. 269, 305–310 (1990).

36. J. J. M. Vitória, D. Trigo, O. A. B. da Cruz e Silva, Revisiting APP secretases: An overview on the holistic effects of retinoic acid receptor stimulation in APP processing. Cell. Mol. Life Sci. 79, 101 (2022).

37. A. Colombo, A. Bastone, C. Ploia, A. Sclip, M. Salmona, G. Forloni, T. Borsello, JNK regulates APP cleavage and degradation in a model of Alzheimer’s disease. Neurobiol. Dis. 33, 518–525 (2009).

38. L. Schnöder, W. Hao, Y. Qin, S. Liu, I. Tomic, X. Liu, K. Fassbender, Y. Liu, Deficiency of neuronal p38α MAPK attenuates amyloid pathology in Alzheimer disease mouse and cell models through facilitating lysosomal degradation of BACE1. J. Biol. Chem. 291, 2067–2079 (2016).

39. J. Brumbaugh, B. Di Stefano, K. Hochedlinger, Reprogramming: Identifying the mechanisms that safeguard cell identity. Development. 146, dev182170 (2019).

40. A. Kuchina, L. Espinar, J. Garcia-Ojalvo, G. M. Süel, Reversible and noisy progression towards a commitment point enables adaptable and reliable cellular decision-making. PLOS Comput. Biol. 7, e1002273 (2011).

41. P. Paganos, J. Ullrich-Lüter, A. Almazán, D. Voronov, J. Carl, A.-C. Zakrzewski, B. Zemann, M. L. Rusciano, T. Sancerni, M. Schauer, O. Akar, F. Caccavale, M. Cocurullo, G. Benvenuto, J. C. Croce, C. Lüter, M. I. Arnone, Single-nucleus profiling highlights the all-brain echinoderm nervous system. Sci. Adv. 11, eadx7753 (2025).

42. Y. Cho, H.-G. Bae, E. Okun, T. V. Arumugam, D.-G. Jo, Physiology and pharmacology of amyloid precursor protein. Pharmacol. Ther. 235, 108122 (2022).

43. G. Merdes, P. Soba, A. Loewer, M. V. Bilic, K. Beyreuther, R. Paro, Interference of human and *Drosophila* APP and APP-like proteins with PNS development in *Drosophila*. EMBO J. 23, 4082–4095 (2004).

44. C. Y. Ewald, C. Li, *Caenorhabditis elegans* as a model organism to study APP function. Exp. Brain Res. 217, 397–411 (2012).

45. M. Nicolas, B. A. Hassan, Amyloid precursor protein and neural development. Development. 141, 2543–2548 (2014).

46. R. E. Diaz, Evo-Devo Path as a bridge between evolution, morphological disparity, and medicine with comments on “hopeful monsters” in the age of genomics. Curr. Mol. Biol. Rep. 6, 79–90 (2020).

47. R. van der Kant, L. S. B. Goldstein, Cellular functions of the amyloid precursor protein from development to dementia. Dev. Cell. 32, 502–515 (2015).

48. J. Dunot, A. Ribera, P. A. Pousinha, H. Marie, Spatiotemporal insights of APP function. Curr. Opin. Neurobiol. 82, 102754 (2023).

49. M. S. Uddin, M. T. Kabir, M. Jalouli, M. A. Rahman, P. Jeandet, T. Behl, A. Alexiou, G. M. Albadrani, M. M. Abdel-Daim, A. Perveen, G. M. Ashraf, Neuroinflammatory signaling in the pathogenesis of Alzheimer’s disease. Curr. Neuropharmacol. 20, 126–146 (2022).

50. S. S. Dominy, C. Lynch, F. Ermini, M. Benedyk, A. Marczyk, A. Konradi, M. Nguyen, U. Haditsch, D. Raha, C. Griffin, L. J. Holsinger, S. Arastu-Kapur, S. Kaba, A. Lee, M. I. Ryder, B. Potempa, P. Mydel, A. Hellvard, K. Adamowicz, H. Hasturk, G. D. Walker, E. C. Reynolds, R. L. M. Faull, M. A. Curtis, M. Dragunow, J. Potempa, *Porphyromonas gingivalis* in Alzheimer’s disease brains: Evidence for disease causation and treatment with small-molecule inhibitors. Sci. Adv. 5, eaau3333 (2019).

51. M. I. Ryder, *Porphyromonas gingivalis* and Alzheimer disease: Recent findings and potential therapies. J. Periodontol. 91(suppl. 1), S45–49 (2020).

52. Z. Huang, M. Hao, N. Shi, X. Wang, L. Yuan, H. Yuan, X. Wang, *Porphyromonas gingivalis*: A potential trigger of neurodegenerative disease. Front. Immunol. 16, 1482033 (2025).

53. M. Dan-Sohkawa, A ‘normal’ development of denuded eggs of the starfish, *Asterina pectinifera*. Dev. Growth Differ. 18, 439–445 (1976).

54. R. Kajitani, D. Yoshimura, M. Okuno, Y. Minakuchi, H. Kagoshima, A. Fujiyama, K. Kubokawa, Y. Kohara, A. Toyoda, T. Itoh, Platanus-allee is a de novo haplotype assembler enabling a comprehensive access to divergent heterozygous regions. Nat. Commun. 10, 1702 (2019).

55. Y. Zhou, B. Zhou, L. Pache, M. Chang, A. H. Khodabakhshi, O. Tanaseichuk, C. Benner, S. K. Chanda, Metascape provides a biologist-oriented resource for the analysis of systems-level datasets. Nat. Commun. 10, 1523 (2019).

