## Supplementary Information for "An APP-centered molecular gateway integrates innate immunity and retinoic acid signaling to drive irreversible metamorphic commitment"

### Supplementary Methods

#### MEGENA implementation

Gene co-expression networks were constructed using the MEGENA R package (v1.3.7). Before analysis, lowly expressed genes were filtered out, and the edgeR-identified set of differentially expressed genes (DEGs) was used as the input. Two preprocessing strategies were evaluated: raw counts (untransformed read counts) and normalized counts **(**log(TMM × CPM + 1) values). In both cases, Pearson correlation coefficients were computed, and significant edges were retained through planar filtering (permutation-based FDR < 0.05). MEGENA was used to construct a hierarchical network and detect modules at multiple scales. Eigengenes were extracted from each module using principal component analysis (prcomp in R) and were defined as the first principal component of standardized gene expression.

#### Dynamic network module (DNM) score calculation

We implemented DNM analysis in R (v4.5.0) based on the principles of Dynamic Network Biomarker (DNB) theory, adapting it to the module eigengenes derived from MEGENA. Module-level applications leverage the stability of the eigengene signal to identify critical transitions in the regulatory architecture, minimizing the noise associated with individual gene fluctuations in low-replicate datasets (as discussed in the Materials and Methods).

The full analysis workflow, including data processing, coefficient of variation (CV) calculations, correlation analyses, and final integration into the DNB scores, is described here.

For each module *m* at time point *t*, the following metrics were computed:

1. **Fluctuation amplitude (ΔCV)**

   We calculated the Coefficient of variation (CV) of the module eigengene *m* at each time point *t*:

   $$CV_{m,t} = \sigma_{m,t} \; / \; |\mu_{m,t}|$$

   where $\mu$ and σ are the mean and standard deviation across replicates. The amplitude ΔCV is the maximum minus the minimum CV across all time points.

2. **Internal correlation**

The average absolute pairwise correlation of eigengenes between module *m* and its sibling modules (sharing the same parent). DNB theory predicts that this correlation increases as the system approaches a critical state (synchronization).

3. **External correlation**
   The average absolute correlation of eigengenes between module *m* and all other modules outside its parent cluster. DNB theory predicts that this correlation decreases as the system separates from the broader network (divergence).

The primary DNM score was defined as the sum of the standardized ($Z$-score) values of the three metrics, where the external correlation term was inverted to reflect divergence.

$$DNM_{score} = Z(\Delta CV) + Z(internal\ correlation) - Z(external\ correlation)$$

**Extended formulation (optional metric):**
An extended formulation ($DNM_{score\_optional}$) was calculated for robustness testing by incorporating a fourth standardized metric, **Isolation from Parent** (1 – |correlation($eigengene_m$, $eigengene_{parent}$)|**).** However, the conclusions in the main text were based on the primary DNM score (integrating the three core metrics).

As no existing package provides this exact functionality, all calculations were performed *de novo* in R. For reproducibility, the final DNM scores and the entire analysis workflow, including the R script (DNM_Analysis_Script.R) and all input data (eigengene_long.csv and module_hierarchy.csv), were provided in a single compressed archive (DNM_Analysis_files. Zip).

### Supplementary figures

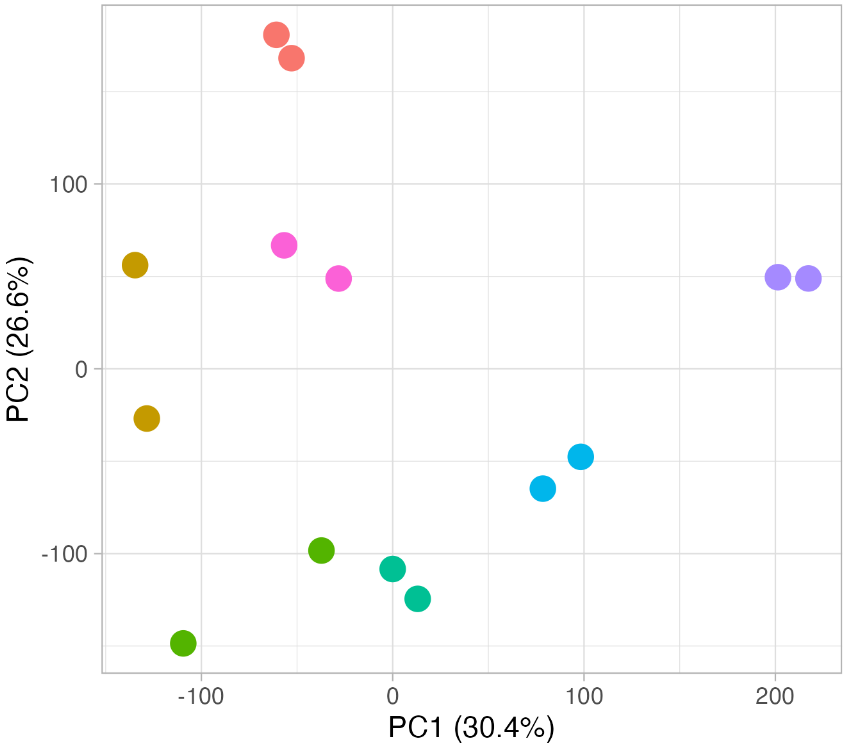


**Figure S1. PCA of transcriptomes across metamorphic stages.**
Principal component analysis (PCA) was performed on RNA-seq transcriptomes from the initial brachiolaria stage and all subsequent metamorphic stages (early, middle, late, prejuvenile, and juvenile), including the retinoic acid (RA) chemically induced early stage (RA-early). Before PCA, transcripts with low read counts were removed, and the data were transformed using the $\log_{10}$(TPM + 1) method. The resulting PC1/PC2 scatter plot showed that biological replicates clustered tightly by their defined morphological stages, confirming that the staging system captured the major sequential transcriptional transitions driving metamorphosis. The first two principal components, PC1 (30.4%) and PC2 (26.6%), accounted for most of the the dataset's variability. Colors represent the metamorphic stages: brachiolaria (red), early (yellow-brown), RA-early (pink), middle (green), late (teal), pre-juvenile (cyan), and juvenile (lavender).

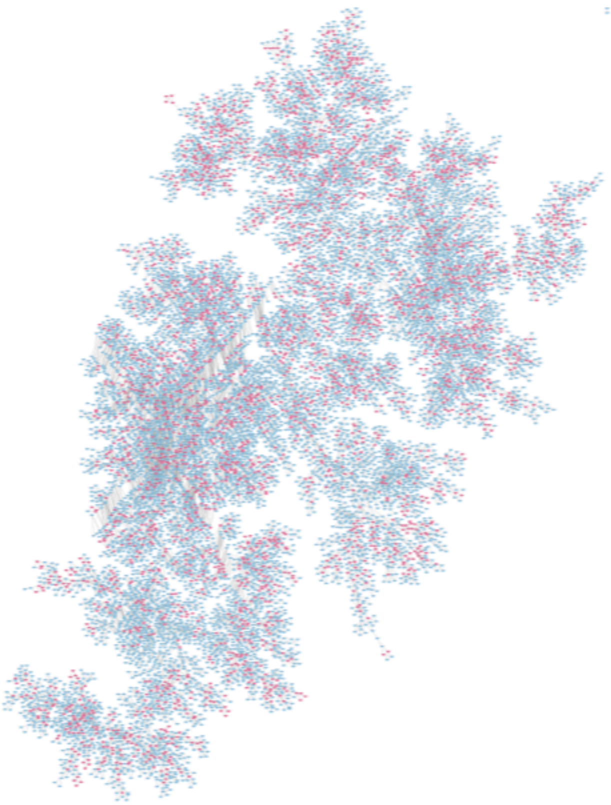

**Figure S2. MEGENA network constructed using normalized log(CPM) expression values.**

The global co-expression network was generated using the Multiscale Embedded Gene Co-expression Network Analysis (MEGENA) method using $\log_{10}(CPM + 1)$-transformed expression values as the input. The network comprises differentially expressed genes (DEGs) across the metamorphic time course (nodes colored light blue), with highly connected hub genes denoted in magenta. As detailed in the Materials and Methods section of the main text, a network constructed from raw read counts was adopted for subsequent analyses, including Dynamic Network Module (DNM) identification. This decision was based on the finding that the raw-count network exhibited stronger functional congruence and superior interpretive power with early-stage comparative and functional data than the network built from log-normalized (CPM) values. The results of functional enrichment analysis performed using the hub genes identified in this log(CPM)-based network are presented in Supplementary Table 2.

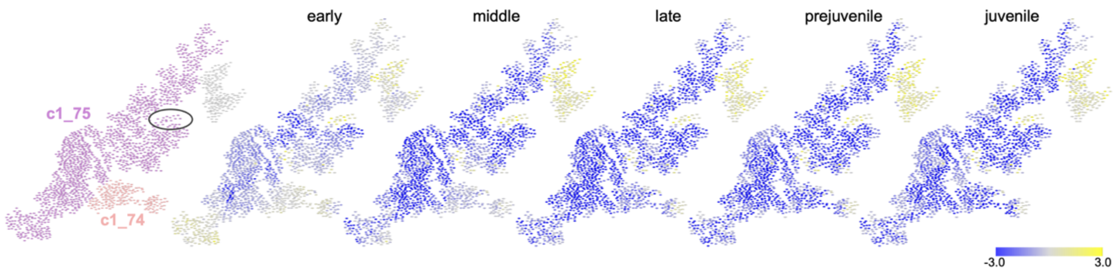


**Figure S3. Expression dynamics of the APP-hub sub-module (c1_75) within the c1_5 module.**

The six panels illustrate the co-expression network structure and expression dynamics of the MEGENA module c1_5 across metamorphosis, providing evidence for the role of APP in signal focusing. The leftmost panel displays the hierarchical structure of c1_5, showing its component submodule c1_75 (purple) and neighboring submodule c1_74 (pink). The APP-containing region within c1_75 that exhibited distinct expression dynamics is demarcated by a black ellipse. The subsequent five panels map the log(counts + 1) expression values of the c1_5 constituent genes onto the network structure at five sequential metamorphic stages: early, middle, late, prejuvenile, and juvenile. Gene expression levels were visualized using a color gradient from low (blue) to high (yellow) with the corresponding color bar indicated in the figure. A comparison across the stages revealed a stage-dependent mass shutdown (decreased expression) for most genes in c1_5. However, a subset of genes within the main c1_5 module, which did not belong to c1_74 or c1_75, maintained relatively high expression, suggesting an independent general maintenance function. In contrast, the APP-associated region of c1_75 demonstrated signal focusing by maintaining consistently high expression throughout metamorphic progression, reinforcing its function as an irreversible commitment gateway.
