## Supplementary Dataset for "An APP-centered molecular gateway integrates innate immunity and retinoic acid signaling to drive irreversible metamorphic commitment": DNM_Analysis_Script.R

```
# ==============================================================================
# DNM_Analysis_Script.R: Dynamic Network Module (DNM) Score Calculation
#
# FIX: Restructured the final 'mutate' block to explicitly name all Z-score 
#      components (e.g., Fluctuation_Zscore) before calculating the final score.
# Logic: Synchronization/Over-coordination (High Internal Corr = High Score)
# ==============================================================================

# Load required libraries
if (!requireNamespace("tidyverse", quietly = TRUE)) {
  install.packages("tidyverse")
}
library(tidyverse)

# --- 1. Data Loading and Preparation (Requires eigengene_long.csv and module_hierarchy.csv) ---

message("--- 1. Loading data and preparation ---")

# Placeholder loading (Replace with your actual data loading)
# NOTE: Assume 'eigengene_long.csv' and 'module_hierarchy.csv' are in the working directory.
eigengene.long <- tryCatch(read_csv("eigengene_long.csv"), error = function(e) {
  stop("Error loading eigengene_long.csv: Please ensure the file exists and is correctly formatted.")
}) %>%
  # Ensure column types are correct
  mutate(module = as.character(module), sample = as.character(sample), timepoint = as.character(timepoint)) 
module_hierarchy <- tryCatch(read_csv("module_hierarchy.csv"), error = function(e) {
  stop("Error loading module_hierarchy.csv: Please ensure the file exists and is correctly formatted.")
}) %>%
  mutate(module = as.character(module), module.parent = as.character(module.parent)) %>%
  filter(!is.na(module.parent))

# Ensure long data has parent info for isolation calculation
if (!("module.parent" %in% names(eigengene.long))) {
  eigengene.long <- eigengene.long %>% left_join(module_hierarchy, by = "module")
}
# Create wide format dataframe for correlation matrix calculation
eigengene.df <- eigengene.long %>%
  select(module, sample, eigengene) %>%
  pivot_wider(names_from = module, values_from = eigengene)

# --- 2. Function Definitions ---

# Function to calculate Isolation from Parent (1 - |r_parent_child|)
calculate_isolation_from_parent <- function(module_name, eig_df_long) {
  eig_df_long <- as.data.frame(eig_df_long)
  
  mod_sub <- eig_df_long %>% filter(module == module_name)
  parent_name <- mod_sub$module.parent[1]
  
  if (is.na(parent_name) || length(parent_name) == 0) return(NA_real_)
  
  parent_sub <- eig_df_long %>%
    filter(module == parent_name, sample %in% mod_sub$sample)
  
  if (nrow(parent_sub) == 0) return(NA_real_)
  
  merged <- inner_join(mod_sub, parent_sub, by = "sample", suffix = c("_child", "_parent"))
  
  cor_val <- suppressWarnings(cor(merged$eigengene_child, merged$eigengene_parent, use = "pairwise.complete.obs"))
  
  if (is.na(cor_val)) return(NA_real_)
  
  return(1 - abs(cor_val))
}

# DNM score calculation function (main logic)
compute_DNM_score <- function(long_eigen, eigengene_df) {
  
  message("Step 1: Compute CV table (Fluctuation Amplitude) ...")
  cv_table <- long_eigen %>%
    group_by(module, timepoint) %>%
    summarise(
      mean_eig = mean(eigengene, na.rm = TRUE),
      sd_eig = sd(eigengene, na.rm = TRUE),
      cv = sd_eig / abs(mean_eig),
      .groups = "drop"
    )
  
  message("Step 2: Summarize CV values ...")
  cv_summary <- cv_table %>%
    group_by(module) %>%
    summarise(
      max_cv = max(cv, na.rm = TRUE),
      min_cv = min(cv, na.rm = TRUE),
      delta_cv = max_cv - min_cv,
      mean_cv = mean(cv, na.rm = TRUE),
      .groups = "drop"
    )
  
  message("Step 3: Compute internal/external correlations (and Z-scores) ...")
  eig_mat <- eigengene.df %>%
    select(-sample) %>%
    select_if(is.numeric) %>%
    as.matrix()
  
  cor_mat <- cor(eig_mat, use = "pairwise.complete.obs")
  
  parent_map <- module_hierarchy %>% filter(!is.na(module.parent))
  
  # Internal Correlation (Synchronization): High correlation = High risk
  internal_corr <- sapply(colnames(cor_mat), function(mod) {
    parent <- parent_map$module.parent[parent_map$module == mod]
    
    # FIX: Handle top-level modules where parent is not found in the map
    if (length(parent) == 0) return(NA)
    
    siblings <- parent_map %>%
      filter(module.parent == parent) %>%
      pull(module)
    siblings <- setdiff(siblings, mod)
    if (length(siblings) == 0 || !all(siblings %in% colnames(cor_mat))) return(NA)
    mean(abs(cor_mat[mod, siblings]), na.rm = TRUE)
  })
  
  # External Correlation (Divergence): Low correlation = High risk
  external_corr <- sapply(colnames(cor_mat), function(mod) {
    parent <- parent_map$module.parent[parent_map$module == mod]
    
    # FIX: Handle top-level modules where parent is not found in the map
    if (length(parent) == 0) {
        # Top-level modules: External correlation is mean corr with ALL others.
        others <- setdiff(colnames(cor_mat), mod)
        if (length(others) == 0) return(NA)
        return(mean(abs(cor_mat[mod, others]), na.rm = TRUE))
    }
    
    # For non-top-level modules, exclude siblings and parent
    exclude <- c(mod, parent_map$module[parent_map$module.parent == parent])
    others <- setdiff(colnames(cor_mat), exclude)
    if (length(others) == 0) return(NA)
    mean(abs(cor_mat[mod, others]), na.rm = TRUE)
  })
  
  cor_stats <- data.frame(
    module = colnames(cor_mat),
    internal_corr = internal_corr,
    external_corr = external_corr
  ) %>%
    mutate(
      # Synchronization Logic (Internal Correlation UP = Score UP)
      InternalSynchronization_Zscore = scale(internal_corr)[,1], 
      # Divergence Logic (External Correlation DOWN = Score UP)
      ExternalDivergence_Zscore = -scale(external_corr)[,1] 
    )
  
  message("Step 4: Local isolation from parent (optional) ...")
  parent_isolation <- sapply(cv_summary$module, calculate_isolation_from_parent, eig_df_long = long_eigen)
  
  message("Step 5: Integrate into final DNM score ...")
  dnb_score_table <- cv_summary %>%
    left_join(cor_stats, by = "module") %>% # Includes InternalSynchronization_Zscore and ExternalDivergence_Zscore
    left_join(module_hierarchy, by = "module") %>%
    mutate(
      isolation_from_parent = parent_isolation,
      
      # 1. Calculate the remaining Z-scores explicitly and name them
      Fluctuation_Zscore = scale(delta_cv)[,1],
      Isolation_from_Parent_Zscore = scale(isolation_from_parent)[,1]
    ) %>%
    # 2. Calculate Final DNM Scores using the named components
    mutate(
      # Final DNM Score: Fluctuation (+) + Internal Synchronization (+) + External Divergence (+)
      DNM_score = Fluctuation_Zscore + InternalSynchronization_Zscore + ExternalDivergence_Zscore,
      DNM_score_optional = DNM_score + Isolation_from_Parent_Zscore
    ) %>%
    arrange(desc(DNM_score))
  
  return(list(
    score_table = dnb_score_table,
    cv_table = cv_table,
    cor_matrix = cor_mat
  ))
}

# --- 4. Execution and Result Output ---
message("--- 4. Computing DNM scores and writing files ---")

# Execute main calculation
result <- compute_DNM_score(eigengene.long, eigengene.df)

# Write out DNM score results
write.table(result$score_table,"DNM_score_table.txt",
            row.names = F, quote = F, sep = "\t")

# Summarize total DNM score by module.parent
parent_summary <- result$score_table %>%
  filter(!is.na(module.parent)) %>%
  group_by(module.parent) %>%
  summarise(total_DNM_score = sum(DNM_score, na.rm = TRUE)) %>%
  arrange(desc(total_DNM_score))
write.table(parent_summary, "parent_DNM_score.txt",
            row.names = F, quote = F, sep = "\t")

message("--- Complete ---")
message("DNM_score_table.txt and parent_DNM_score.txt have been generated.")
```
